# Dissection of Polycomb complexes reveals canonical PRC1-independent 3D chromatin interactions

**DOI:** 10.64898/2026.10.13.758654

**Authors:** Miles K. Huseyin, Sumin Kim, Jack Toppen, Neil P. Blackledge, Anders S. Hansen

## Abstract

The Polycomb system is an essential regulator of developmental genes which controls chromatin state through histone modifications and 3D chromatin organization. Although Polycomb-mediated histone modification is well understood, how Polycomb-mediated 3D chromatin interactions form remains unclear. To address this gap, we combine comprehensive Polycomb subunit degrons with Region Capture Micro-C, ChIP-seq, nascent transcriptomics, and live-cell imaging of DNA loci and Polycomb bodies. We find that, within canonical Polycomb repressive complex 1 (cPRC1), polyhomeotic proteins are the primary contributors to interactions. In surprising contrast to existing models, we find that Polycomb-mediated interactions persist in the absence of cPRC1, and combined depletion of cPRC1 with individual variant PRC1 and PRC2 complexes indicates that the remaining interactions do not depend on a single complex. These results raise the previously unexplored possibility of a more general mechanism for Polycomb-mediated interactions not requiring cPRC1.

## INTRODUCTION

Chromatin represents the key medium for the regulation of gene expression in eukaryotes^1^. Transcriptional regulators can manipulate chromatin through a multitude of mechanisms, including chemical modification of DNA and histone proteins^2^, controlling the accessibility of DNA^3^, and mediating chromatin interactions in 3D space^4^. Distinguishing the causes and consequences of individual regulatory mechanisms has proven difficult as perturbation of regulatory proteins often impacts multiple chromatin features simultaneously^5,6^. The Polycomb repressive system, an essential repressor of developmental genes, is a paradigm of such multi-layered regulation, being known to deposit histone modifications^7,8^, compact chromatin^9–13^, and mediate 3D chromatin interactions^11,14–22^.

The Polycomb system consists of two main complexes, Polycomb repressive complexes 1 and 2 (PRC1 and PRC2), with distinct composition and functions. PRC1 catalyzes deposition of monoubiquitylation on lysine 119 of histone H2A (H2AK119ub1)^8,23^, while PRC2 has methyltransferase activity specific to lysine 27 of histone H3 (H3K27me1/2/3)^7,24–26^. PRC1 complexes form around a heterodimer of RING1A or RING1B and one of six PCGF proteins (PCGF1/2/3/4/5/6)^27–30^, with complexes differing in catalytic activity and subunit composition based on the incorporated PCGF protein^31–39^. PRC1 complexes are further broadly categorized into canonical (cPRC1) and variant (vPRC1, or non-canonical) complexes, based on their subunit composition^27,34^. Similarly, PRC2 complexes consist of a conserved core^40,29,30^, along with variable auxiliary subunits which define the catalytic activity and chromatin binding of the complex^41,42^. Polycomb complexes exhibit interdependency in chromatin occupancy: a subset of PRC2 complexes bind to H2AK119ub1 deposited by vPRC1, while cPRC1 binds to chromatin through either recognition of H3K27me3 or PRC2 itself^43–47,7,25,48–55^. This compositional complexity and interplay between the complexes have made the Polycomb system difficult to study.

Beyond histone-modifying activity, the Polycomb system also drives chromatin compaction and 3D chromatin interactions^56^. These functions are thought to be driven by cPRC1 complexes which contain PCGF2/4 in combination with chromobox (CBX) and polyhomeotic (PHC) subunits^27,29,30,34^. Consistent with this, tethering of PCGF2 is sufficient to drive interactions between the tethering locus and Polycomb sites^57^. Both the CBX and PHC proteins can self-interact, and these subunits are proposed to be directly responsible for chromatin compaction and Polycomb-mediated 3D chromatin interactions (henceforth “Polycomb-mediated interactions”)^16,11,58,9,10,12,59–61,22,62–64^. Polycomb proteins also form nuclear foci known as Polycomb bodies, which exhibit higher enrichment of cPRC1 subunits than vPRC1 subunits^65^. Microscopy experiments have also shown colocalization of Polycomb bodies with Polycomb-repressed genes, suggesting that Polycomb bodies may represent interacting Polycomb-occupied loci^16,66–68,63^. However, the function of Polycomb-mediated interactions remains unclear^56^. Previous work in which Polycomb-mediated interactions were indirectly perturbed observed changes in transcription of Polycomb-repressed genes^21,69^. Despite this, cPRC1 has substantially lower ubiquitin E3 ligase activity than vPRC1 both in vitro and in cells, and makes only minimal contributions to gene repression^27,36,37,70^. This emphasizes a key open question: if disrupting Polycomb-mediated interactions affects transcription^21,69^ and cPRC1 is solely responsible for Polycomb-mediated interactions^11,16,58,62,63^, then why does cPRC1 not affect transcription?

Although the roles of Polycomb-mediated histone modifications have begun to be understood, how Polycomb contributes to 3D genome organization remains unclear. Perturbing total PRC1 or PRC2 not only disrupts Polycomb-mediated interactions, but also impacts the Polycomb histone modifications, affects other Polycomb complexes through feedback between PRC1 and PRC2, and can lead to substantial gene expression changes^9,15,17,21,63,71,72^. Therefore, although multiple studies have examined the effect on Polycomb-mediated interactions with PRC1 perturbation, it is difficult to conclude whether loss of 3D interactions follows directly from PRC1 perturbation or is the consequence of downstream effects^9,17,21,63,71,72^. To date, studies have addressed this complexity by deleting or mutating PHC proteins^11,16^ or conditionally depleting core cPRC1 subunits^57,73^. These perturbations weaken interactions between Polycomb-occupied loci, lead to decompaction of individual loci, and disrupt Polycomb body formation^11,16,57,73^. However, these studies have largely relied on long-term deletion or mutation of individual subunits, or ectopic expression, limiting their ability to comprehensively address how the individual subunits work together in endogenous contexts. Furthermore, most genomics studies of Polycomb-mediated interactions have assessed 3D genome organization through Hi-C-based approaches^11,15,17,21,63^. These methods are limited in their ability to identify and quantify weak interactions compared to newer approaches, such as Micro-C^74,75^. Besides genomics, Polycomb-mediated interactions have been studied using microscopy methods based on denaturing fluorescence in situ hybridization (FISH)^9,11,16,63,72,76^. However, these methods suffer from the need to crosslink cells and denature the DNA, processes which have been shown to perturb chromatin structure and nuclear foci^77–79^.

Here, we overcome these limitations using a panel of degron cell lines with Region Capture Micro-C (RCMC)^80,81^ and live-cell imaging to comprehensively study Polycomb-mediated interactions and Polycomb bodies. We first establish a maximal effect size by depleting both PRC1 and PRC2 and then more specifically deplete CBX and PHC proteins to determine their contributions to Polycomb-mediated interactions. We further deplete cPRC1 complexes completely and find that a low level of interaction between Polycomb-occupied sites persists in the absence of intact cPRC1. Lastly, we combine our cPRC1 degron with simultaneous depletion of other Polycomb complexes and find that no combination fully recapitulates interaction losses in a complete PRC1 degron. Together, this suggests the existence of a general mechanism for Polycomb-mediated interactions which does not require cPRC1.

## RESULTS

### PRC1 is necessary for Polycomb-mediated 3D chromatin interactions

To quantify the contributions of different Polycomb subunits to Polycomb-mediated 3D chromatin interactions, we first aimed to establish a condition for maximal interaction loss using inducible depletion. We therefore generated Polycomb degron cell lines in mouse embryonic stem cells (mESCs), where the Polycomb system has been extensively studied and is known to form strong 3D chromatin interactions^15,17,19,74^. Although previous studies have found that PRC1 depletion or deletion leads to a significant reduction in chromatin occupancy of PRC2, some level of PRC2 remains on chromatin^44,70,71,82^. We therefore wanted to test whether acute depletion of PRC1 alone was sufficient to achieve maximal loss of Polycomb-mediated interactions or if both complexes contribute. To this end, we generated PRC1 (PRC1^deg^, RING1A/B) and PRC1/2 (PRC1/2^deg^, RING1A/B SUZ12) degron cell lines with core subunits of each complex tagged using the dTAG inducible degron system^83^ (Fig. 1a and Fig. S1a,b). Both cell lines showed efficient depletion of tagged proteins in 4 hours of treatment (Fig. 1b and Fig. S1c), and near-complete loss of bulk H2AK119ub1 (Fig. S1d). Only the PRC1/2^deg^ cell line showed a moderate decrease in bulk H3K27me3, reduced to ∼80% of untreated (Fig. S1d). This is consistent with prior work showing that H3K27me3 has a long half-life compared to H2AK119ub1, which turns over rapidly (*t*_½_ = 90 minutes)^53,65,82,84^. In the PRC1/2^deg^ cell line, we further confirmed that RING1B, SUZ12, and cPRC1 subunits (PCGF2, PHC1, CBX2, CBX7) were lost from Polycomb-occupied sites by ChIP-seq (Fig. 1c and Fig. S1e,f). Similarly, H2AK119ub1 showed near-complete loss at the same sites, while H3K27me3 was reduced by around 50% (Fig. S1g). Measuring nascent transcription using TT_chem_-seq^85^, we also observed that the gene expression changes in both the PRC1^deg^ and PRC1/2^deg^ broadly recapitulated those seen for a prior PRC1 degron cell line at 4 h^82^ (Fig. S1h,i). Although short-term SUZ12 depletion has been previously shown not to affect gene expression^82^, we noted that PRC1/2^deg^ exhibited slightly increased gene expression changes compared to PRC1^deg^, indicating a possible redundant role for PRC2 (Fig. S1j).

**Figure 1.**
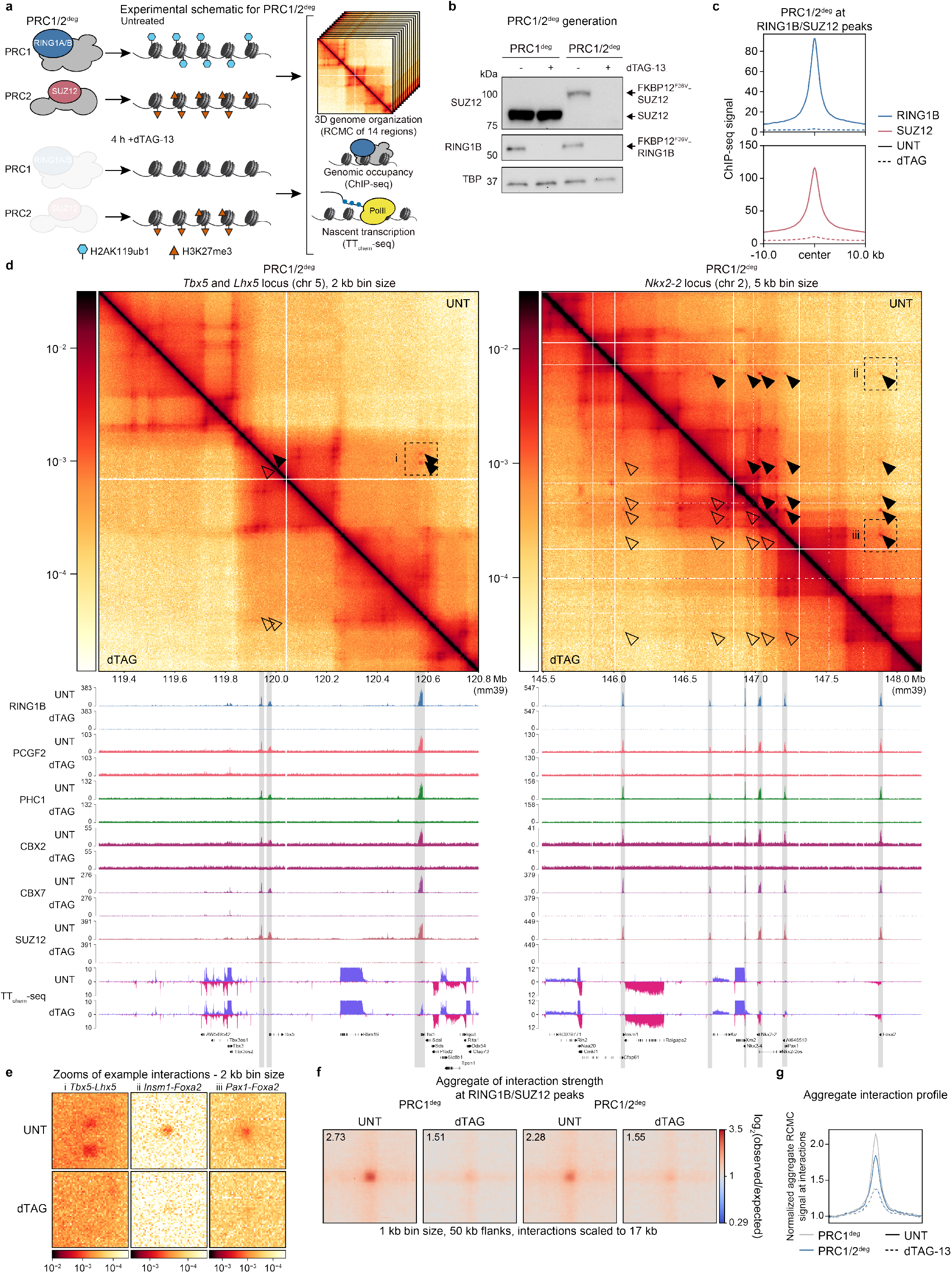
PRC1 drives Polycomb-mediated interactions. **a**, A schematic illustrating the Polycomb complexes and histone modifications in untreated and 4 h dTAG-13 treated PRC1/2^deg^ (RING1A/B SUZ12 degron) cells, and an overview of the genomic protocols performed. **b**, Western blot validation of depletion following 4 h dTAG-13 treatment in PRC1^deg^ and PRC1/2^deg^ cell lines. Depletion of RING1B is shown for both cell lines, tagging and depletion of SUZ12 is shown for PRC1/2^deg^, and TBP is included as a loading control. *n* = 1 biological replicate. **c**, Metaplots showing loss of RING1B and SUZ12 ChIP-seq signal at RING1B/SUZ12 co-occupied peaks following 4 h dTAG-13 treatment in PRC1/2^deg^. Data represent the merge of *n* = 3 biological replicates per condition. **d**, RCMC log_10_-scaled contact maps (upper) with corresponding ChIP-seq and TT_chem_-seq tracks (lower) in PRC1/2^deg^ showing changes in contact frequency, Polycomb genomic occupancy, and gene expression following 4 h dTAG-13 treatment. Filled black arrowheads indicate Polycomb-mediated interactions present in untreated (UNT), outlined arrowheads indicate the same interactions which are lost after PRC1/2 depletion (dTAG). Data shown are merges of *n* = 2 (RCMC) or *n* = 3 (ChIP-seq and TT_chem_-seq) biological replicates. **e**, Zooms of interactions from RCMC contact map regions labeled i, ii, and iii in **d** in untreated and dTAG-13 treatment, labeled according to the genes at the interacting loci. Contact map zooms are plotted at 2 kb bin size. **f**, Aggregate pileups of interactions (log_2_(observed/expected)) between RING1B/SUZ12 co-occupied sites in PRC1^deg^ and PRC1/2^deg^ with or without 4 h dTAG-13 treatment. Sites are scaled to 17 kb with 50 kb flanks, with data plotted at 1 kb bin size. Values in the upper left corner of each plot indicate the mean value of the central 3×3 pixels of the pileup. **g**, A 1D average plot across the 21 middle diagonals of the aggregate pileups in **f** for PRC1^deg^ and PRC1/2^deg^, normalized to the mean of the 10 bins on the left- and right-most sides.

Having validated our degron cell lines, we performed RCMC^80,81^ to investigate their effect on Polycomb-mediated 3D chromatin interactions. RCMC allowed us to generate deeply sequenced contact maps of regions containing Polycomb-mediated interactions in many conditions and without sequencing across the entire genome, providing a sensitive readout of changes in 3D chromatin interaction strength. Using existing 3D genomic data as a reference^74^, we selected 14 genomic regions containing interactions between Polycomb-occupied genes, totaling 25.5 Mb in linear genomic distance (Tables S1 and S2). We were able to achieve ∼100-1000-fold deeper contact maps of Polycomb-mediated interactions compared to a previous study of PRC1 depletion using Hi-C^21^ (Fig. 1d and Fig. S2a,b). This allowed us to observe all of the Polycomb-mediated interactions visible in the Hi-C data, as well as interactions that were considerably noisier or not visible in the Hi-C data, enabling more precise quantification of their interaction strength. Notably, previous work has established that these sites maintain accessibility at similar levels following conditional deletion of PRC1 and PRC2^86^, indicating that changes in interaction strength were due to Polycomb depletion and not changes to accessibility (Fig. S2a).

Using our RCMC data, we determined that dTAG-13 treatment in PRC1/2^deg^ led to large reductions in Polycomb-mediated interaction strength (Fig. 1d,e,f,g and Fig. S3). Although some enrichment of interaction strength remained, these sites have minimal enrichment of non-Polycomb chromatin factors (Fig. S4a). We further categorized interactions based on whether or not they had the major looping factors CTCF and cohesin at both anchors. Although interactions with CTCF/cohesin at both anchors were stronger overall, both before and after depletion, both sets of interactions were strongly decreased following dTAG-13 treatment (Fig. S4b). This suggests that the remaining interaction strength does not arise from CTCF/cohesin. Instead, it is likely due to either incomplete loss of interactions at 4 h, remaining H3K27me3, or weak interactions of derepressed genes following Polycomb depletion. Overall, we found that PRC1^deg^ and PRC1/2^deg^ reduced the aggregate interaction strength between Polycomb sites in the captured regions to similarly low levels (Fig. 1f,g). This suggests that the remaining PRC2 following depletion of only PRC1 does not sustain interactions alone, and that PRC1 depletion is sufficient to maximally weaken Polycomb-mediated interactions after four hours. This therefore establishes a maximum weakening of Polycomb mediated interactions for further comparison.

### PRC2 depletion partly weakens Polycomb-mediated interactions

While PRC2 depletion in addition to PRC1 depletion does not further weaken Polycomb mediated interactions, we wanted to test whether this ruled out PRC2 contributing to Polycomb-mediated interactions. PRC2 is known to support cPRC1 chromatin binding both indirectly, by placing H3K27me3 which is recognized by cPRC1^51,54,50,55,33^, and directly, through supporting CBX7 chromatin occupancy^52,53^, and may thus contribute to Polycomb-mediated interactions. Therefore, we generated a SUZ12-only degron cell line (PRC2^deg^) as previously described^65^ to test the role of PRC2 alone in contributing to 3D chromatin interactions (Fig. S5a). Although SUZ12 depletion on short timescales in mESCs does not lead to changes in gene expression^5^, we observed a noticeable decrease in Polycomb-mediated interaction strength (Fig. S5b,c,d). This effect was weaker than that of PRC1 depletion, suggesting that PRC2 only contributes a fraction of Polycomb-mediated interaction strength (Fig. S5d). The decrease in Polycomb-mediated interaction strength in this condition might arise from a combination of two sources: lost direct contribution of PRC2 to cPRC1 chromatin occupancy and a partial reduction in H3K27me3. However, from this experiment, we cannot exclude the possibility that PRC2 may also directly support interactions via an unknown mechanism.

### PHC proteins are the major driver of Polycomb-mediated interactions in cPRC1

Having established the requirement of PRC1 for Polycomb-mediated interactions, we next sought to determine the relative contributions of individual cPRC1 subunits. Within cPRC1, two major mechanisms contributing to Polycomb-mediated interactions have been proposed: interactions between the Compaction and Phase Separation (CaPS) domain of CBX proteins^58^, and polymerization of the Sterile Alpha Motif (SAM) domain of PHC proteins^11,16^ (Fig. 2a).

**Figure 2.**
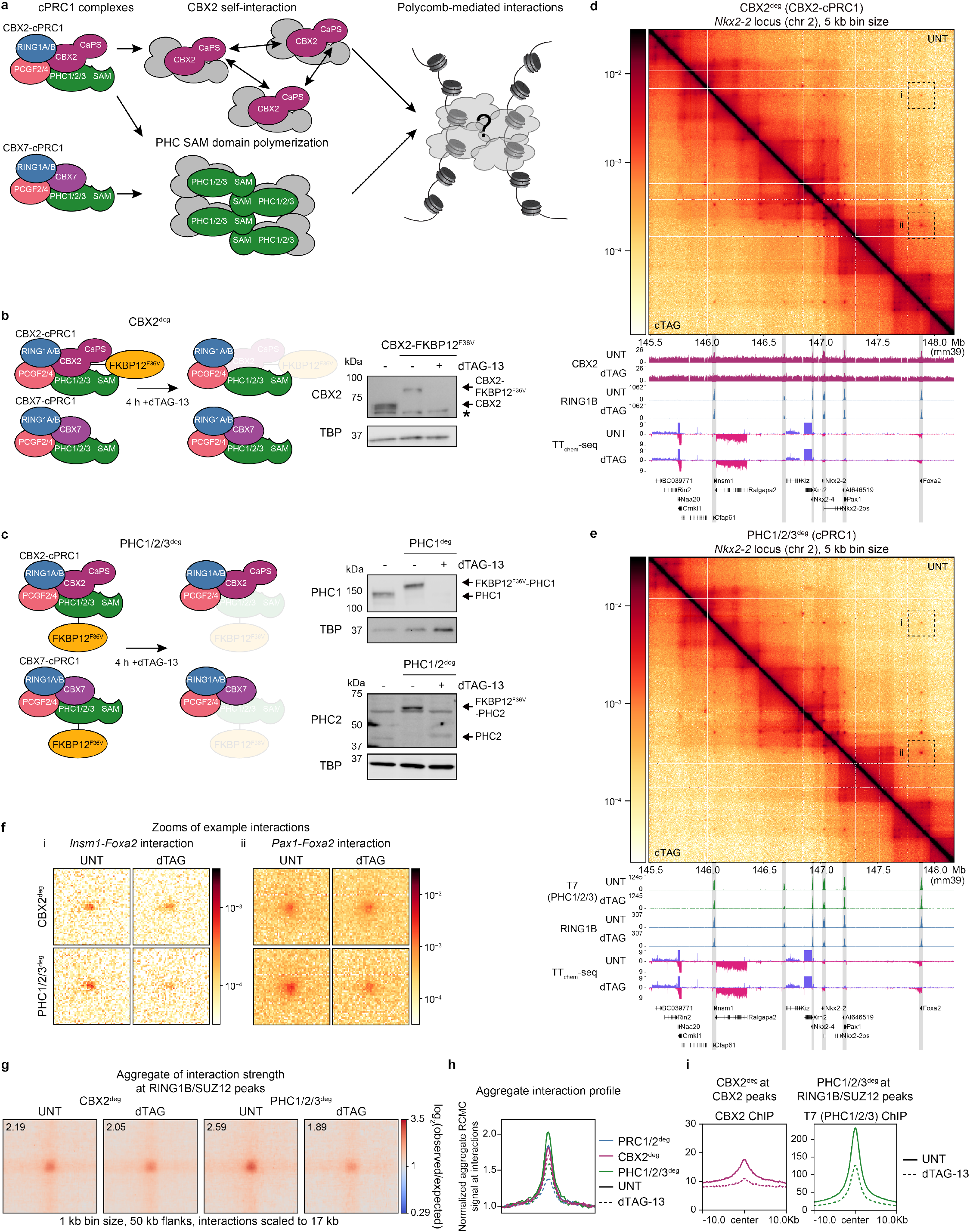
PHC proteins and not CBX2 are required for full Polycomb-mediated interaction strength. **a**, A schematic illustrating cPRC1 complexes in mESCs (left), proposed mechanisms of interaction formation by cPRC1 (middle), via either self-interaction of CBX2 or polymerization of PHC proteins, and how these mechanisms may contribute to Polycomb-mediated interactions (right). **b-c**, A schematic of CBX2 depletion in the CBX2^deg^ (**b**) or PHC1/2/3 in the PHC1/2/3^deg^ cell line (**c**) (left), and western blots showing CBX2 (**b**) or PHC1 and PHC2 (**c**) tagging and depletion following 4 h dTAG-13, including TBP as a loading control. *n* = 1 biological replicate for all blots. **d-e**, RCMC log_10_-scaled contact maps (upper) with corresponding ChIP-seq and TT_chem_-seq tracks (lower) in CBX2^deg^ (**d**) or PHC1/2/3^deg^ (**e**) showing changes in contact frequency, Polycomb genomic occupancy, and gene expression following 4 h dTAG-13 treatment. Data shown are merges of *n* = 2 (RCMC) or *n* = 3 (ChIP-seq and TT_chem_-seq) biological replicates. **f**, Zooms of interactions from RCMC contact map regions labeled i and ii in **d** and **e** in untreated and dTAG-13 treatment, labeled according to the genes at the interacting loci. Contact map zooms are plotted at 2 kb bin size. **g**, Aggregate pileups of interactions (log_2_(observed/expected)) between RING1B/SUZ12 co-occupied sites in CBX2^deg^ and PHC1/2/3^deg^ with or without 4 h dTAG-13 treatment. Sites are scaled to 17 kb with 50 kb flanks, with data plotted at 1 kb bin size. Values in the upper left corner of each plot indicate the mean value of the central 3×3 pixels of the pileup. **h**, A 1D average plot across the 21 middle diagonals of the aggregate pileups in **g** and **Fig. 1f** for PRC1/2^deg^, CBX2^deg^, and PHC1/2/3^deg^, normalized to the mean of the 10 bins on the left- and right-most sides. **i**, Metaplots showing changes in CBX2 ChIP-seq at CBX2 peaks (left) and PHC1/2/3 ChIP-seq signal at RING1B/SUZ12 co-occupied peaks (right) following 4 h dTAG-13 treatment in CBX2^deg^ and PHC1/2/3^deg^. Data represent the merge of *n* = 3 biological replicates per condition.

There are five cPRC1-associated CBX proteins (CBX2, 4, 6, 7, and 8). All have a CaPS domain except CBX7^12^, the major expressed CBX protein in mESCs^33,58^. The CaPS domain-containing CBX2 is known to also be expressed in mESCs, albeit at low levels^58^. Therefore, while CBX2 may drive interactions through its CaPS domain, CBX7 is thought to be the main driver of cPRC1 chromatin occupancy in mESCs as a result of its higher expression. Of the other *Cbx* genes, *Cbx4* and *Cbx8* are minimally expressed in our mESC TT_chem_-seq data, and in fact are Polycomb-repressed (Fig. S6a). We were unable to detect CBX6 at the protein level despite nascent transcription of the gene, consistent with previous work that did not detect CBX6 in PRC1 by mass spectrometry in mESCs^29,33^ (Fig. S6a). *Phc1* predominates the expressed *Phc* genes, with low *Phc2* and *Phc3* expression in TT_chem_-seq data (Fig. S6a). However, peptides from both proteins have previously been reported in mESC cPRC1 complexes^29,33^.

We therefore generated three degrons of cPRC1 subunits: a CBX2 degron (CBX2^deg^) to test the role of the CaPS domain, a PHC1/2/3 triple degron (PHC1/2/3^deg^) to test the role of PHC polymerization, and a CBX7 degron (CBX7^deg^) for the role of general cPRC1 chromatin occupancy (Fig. 2b,c and Fig. S6b). All tagged proteins were detectable in untreated cells and depleted after 4 h of dTAG-13 treatment except PHC3, which we could not detect in unedited or untreated cells (Fig. 2b,c and Fig. S6c). We therefore confirmed PHC3 was tagged at the DNA level but concluded that it may be undetectable at the protein level due to its similar size to PHC1 and potentially low expression (Fig. S6d). Consistent with previous observations that cPRC1 disruption by PCGF2/4 deletion led to only minor changes in transcription^70^, dTAG-13 treatment in CBX2^deg^, CBX7^deg^, and PHC1/2/3^deg^ resulted in no significant gene expression changes (Fig. S6e).

We then performed RCMC for CBX2^deg^ and PHC1/2/3^deg^ and assessed the change in Polycomb-mediated interactions following dTAG-13 treatment (Fig. 2d,e). Consistent with a previous report that CBX2 depletion does not affect interaction strength^58^, dTAG-13 treatment of CBX2^deg^ had a limited effect on these interactions (Fig. 2d,f,g,h). By contrast, dTAG-13 treatment of PHC1/2/3^deg^ broadly weakened Polycomb-mediated interactions (Fig. 2e,f,g,h). However, depletion of PHC1/2/3 did not match the extent of reduction in Polycomb-mediated interactions of the PRC1/2^deg^ cell line. We also observed that, despite the depletion of PHC1 and PHC2 appearing complete in western blotting, T7 ChIP-seq of the tagged PHC proteins showed that around 50% of the ChIP-seq signal remained at Polycomb target sites (Fig. 2e,i). This possibly explains the difference in effect size between PHC1/2/3^deg^ and PRC1/2^deg^. Although PHC proteins were not completely lost in ChIP-seq, RING1B signal was substantially reduced (Fig. S6f). This indicates a disruption to either cPRC1 chromatin binding or complex integrity, or that PHC depletion also targets the remainder of the cPRC1 complex.

Depletion of CBX7 caused strong reductions in chromatin occupancy of both CBX7 and PHC1 in ChIP-seq, with little to no reduction in RING1B occupancy (Fig. S6g). Intriguingly, we also observed a modest decrease in SUZ12 chromatin occupancy (Fig. S6g). It has been suggested previously that PRC2 directly contributes to CBX7 chromatin occupancy^52,53^, and this observation indicates this interaction may be at least partly reciprocal in nature. Despite these occupancy changes, we observed minimal effects on Polycomb-mediated interaction strength (Fig. S6h,i,j). However, the CBX7^deg^ cell line also possessed much weaker Polycomb-mediated interaction strength in untreated cells than other cell lines, comparable to PHC1/2/3^deg^ following dTAG-13 treatment (Fig. 2h and Fig. S6j). CBX7 and PHC1 could still be detected by ChIP on chromatin in untreated cells, and CBX7 depletion was sufficient to displace PHC1 from chromatin (Fig. S6g). This indicates that in untreated CBX7^deg^ cells, tagging did not perturb CBX7 binding or cPRC1 integrity. The already weakened Polycomb interactions may therefore arise from clonal differences in Polycomb-mediated interaction strength. Despite having weaker untreated interactions, CBX7^deg^ also did not recapitulate the interaction strength decrease of PRC1/2^deg^ before or after dTAG-13 treatment (Fig. S6j).

Together, these results suggest that the primary driver within cPRC1 of Polycomb-mediated interactions in mESCs is PHC proteins, while self-interaction by CBX2 plays a minimal role. However, PHC1/2/3 depletion did not match the Polycomb-mediated interaction loss which we had observed in the PRC1/2^deg^ cell line, potentially due to incomplete loss of PHC proteins from chromatin.

### Polycomb-mediated interactions persist in the absence of cPRC1

In order to fully remove PHC proteins from chromatin without perturbing total PRC1, we aimed to disrupt cPRC1 complexes as a whole. To this end, we generated two further degron cell lines. We first generated a double PCGF2/4^deg^ cell line in which core cPRC1 subunits PCGF2 and PCGF4 can be degraded (Fig. 3a,b and Fig. S7a, b). Although the PHC1/2/3^deg^ cell line did not show complete loss of PHC chromatin occupancy in ChIP-seq, we saw complete loss of PHC1 ChIP-seq signal in PCGF2/4^deg^ cells (Fig. 3c). Having also determined that CBX7 depletion is sufficient to disrupt PHC1 chromatin occupancy, we generated a quintuple PHC1/2/3 CBX2/7^deg^ cell line to deplete all cPRC1 auxiliary subunits simultaneously (Fig. S7c). Cellular fractionation in the PHC1/2/3 CBX2/7^deg^ cell line showed that tagged proteins were lost in all fractions, and that extended depletion times up to 48 h did not lead to further reductions in protein level (Fig. S7d). Based on these results, we used the PCGF2/4^deg^ and PHC1/2/3 CBX2/7^deg^ as exhaustive depletions of cPRC1.

**Figure 3.**
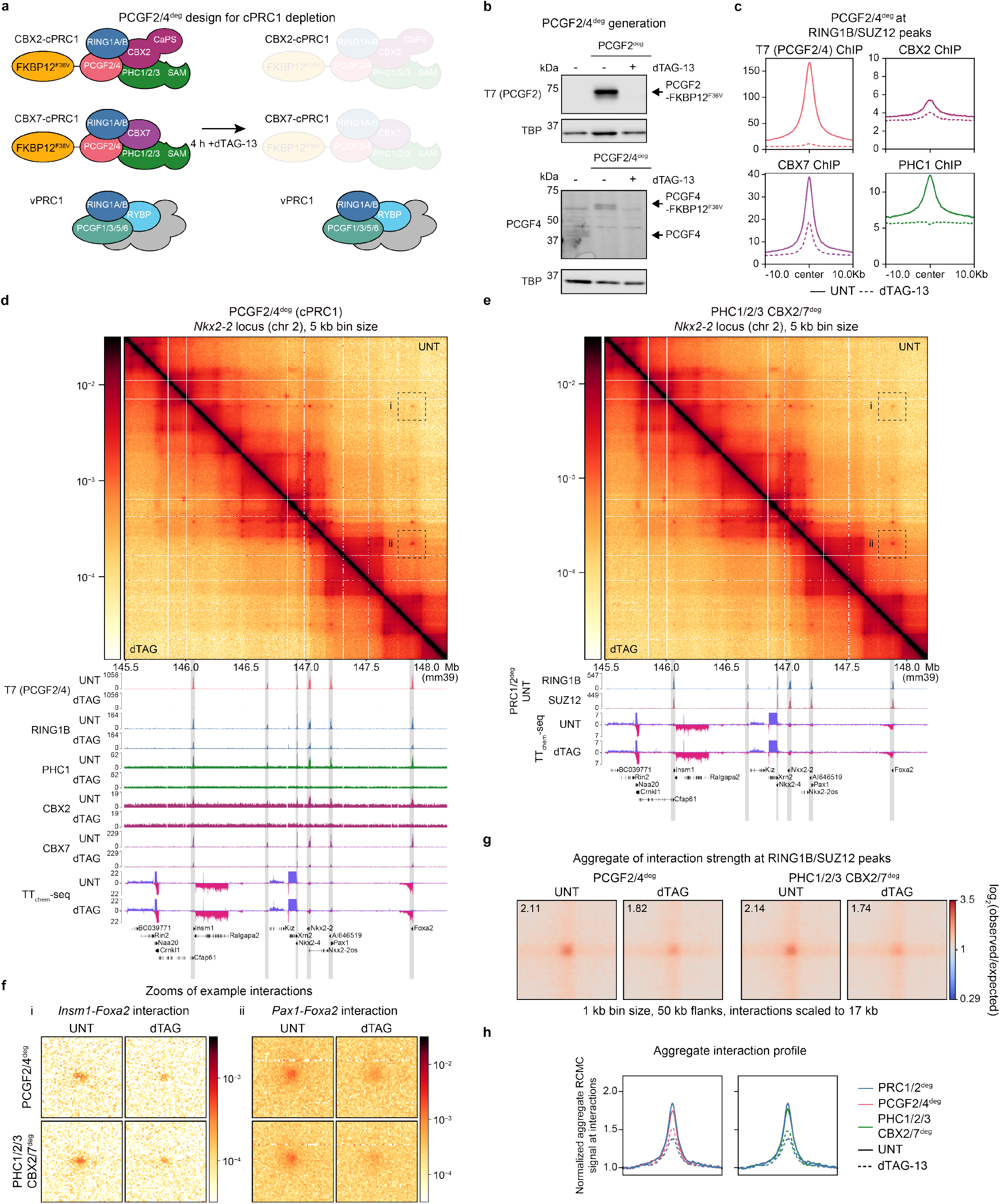
Disruption of cPRC1 weakens but does not ablate Polycomb-mediated interactions. **a**, A schematic illustrating PRC1 complexes in mESCs split between cPRC1 and vPRC1 complexes, and the anticipated effect of depletion of core cPRC1 subunits PCGF2/4. **b**, Western blots showing tagging and depletion of PCGF2 and PCGF4. PCGF2 is probed for with an antibody to the T7 epitope tag included with the degron tag due to the lack of an antibody usable for western blotting. TBP is included as a loading control. *n* = 1 biological replicate for both blots. **c**, Metaplots showing loss of ChIP-seq signal for T7, representing PCGF2/4, and cPRC1 subunits CBX2, CBX7, and PHC1 at RING1B/SUZ12 co-occupied peaks following 4 h dTAG-13 treatment in PCGF2/4^deg^. Data represent the merge of *n* = 3 biological replicates per condition. **d-e**, RCMC log_10_-scaled contact maps (upper) with corresponding ChIP-seq and TT_chem_-seq tracks (lower) in PCGF2/4^deg^ (**d**) and PHC1/2/3 CBX2/7^deg^ (**e**) showing changes in contact frequency, Polycomb genomic occupancy, and gene expression following 4 h dTAG-13 treatment. Data shown are merges of *n* = 2 (RCMC) or *n* = 3 (ChIP-seq and TT_chem_-seq) biological replicates. **f**, Zooms of interactions from RCMC contact map regions labeled i and ii in **d** and **e** in untreated and dTAG-13 treatment, labeled according to the genes at the interacting loci. Contact map zooms are plotted at 2 kb bin size. **g**, Aggregate pileups of interactions (log_2_(observed/expected)) between RING1B/SUZ12 co-occupied sites in PCGF2/4^deg^ and PHC1/2/3 CBX2/7^deg^ with or without 4 h dTAG-13 treatment. Sites are scaled to 17 kb with 50 kb flanks, with data plotted at 1 kb bin size. Values in the upper left corner of each plot indicate the mean value of the central 3×3 pixels of the pileup. **h**, 1D average plots across the 21 middle diagonals of the aggregate pileups in **g** and **Fig. 1f** for PRC1/2^deg^ and PCGF2/4^deg^, normalized to the mean of the 10 bins on the left- and right-most sides.

In line with a previous PCGF2/4 conditional deletion^70^, a small number of genes (52) were significantly derepressed following PCGF2/4 depletion, while most genes remained unchanged (Fig. S7e). Conversely, in PHC1/2/3 CBX2/7^deg^, we observed no differentially expressed genes, consistent with our individual depletions of these proteins (Fig. S7f). This difference may point to a role for catalytically active PCGF2/4-containing vPRC1 complexes^27,37^ at a small number of genes, independent of cPRC1. Overall, these results are consistent with a minimal repressive role for cPRC1 complexes in mESCs^70^.

We then performed RCMC on the PCGF2/4^deg^ and PHC1/2/3 CBX2/7^deg^ cell lines. In both cell lines, dTAG-13 treatment caused a strong reduction in Polycomb-mediated interaction strength, comparable to that of the PHC1/2/3^deg^ cell line, but did not match the effect size in PRC1/2^deg^ (Fig. 3d,e,f,g,h). This result was surprising, given the prevailing view that cPRC1 is the sole driver of these interactions^11,58,62,63^, and suggests the existence of a cPRC1-independent mechanism for Polycomb-mediated interactions.

### Super-resolution locus imaging of Polycomb loci recapitulates RCMC interaction changes

Having found that depletion of cPRC1 was not equivalent to full PRC1 depletion in RCMC, we wanted to validate this observation using a microscopy-based approach. Previous experiments examining the role of Polycomb proteins in mediating 3D genome interactions have measured distances between Polycomb-occupied loci using FISH and related microscopy techniques^9,11,16,62,76^. However, these methods generally use fixation and denaturing of DNA, which can perturb nuclear organization^77–79^. We therefore used super-resolution live-cell microscopy to quantify the 3D distances between two Polycomb-repressed genes. The *Tbx5* and *Lhx5* genes are separated by ∼600 kb and form strong interactions in our RCMC data (Fig. 1d). We inserted previously validated ParB/ParS labels for live-cell imaging^87,88^ upstream of the promoters of each gene in the PRC1/2^deg^,PHC1/2/3^deg^, and PCGF2/4^deg^ lines (Fig. 4a,b). As a negative control for a non-interacting pair that would not be expected to be affected by Polycomb, we also generated cell lines with the same *Tbx5* insertion and an array inserted at a non-Polycomb-occupied intergenic region ∼600 kb upstream (Fig. 4a).

**Figure 4.**
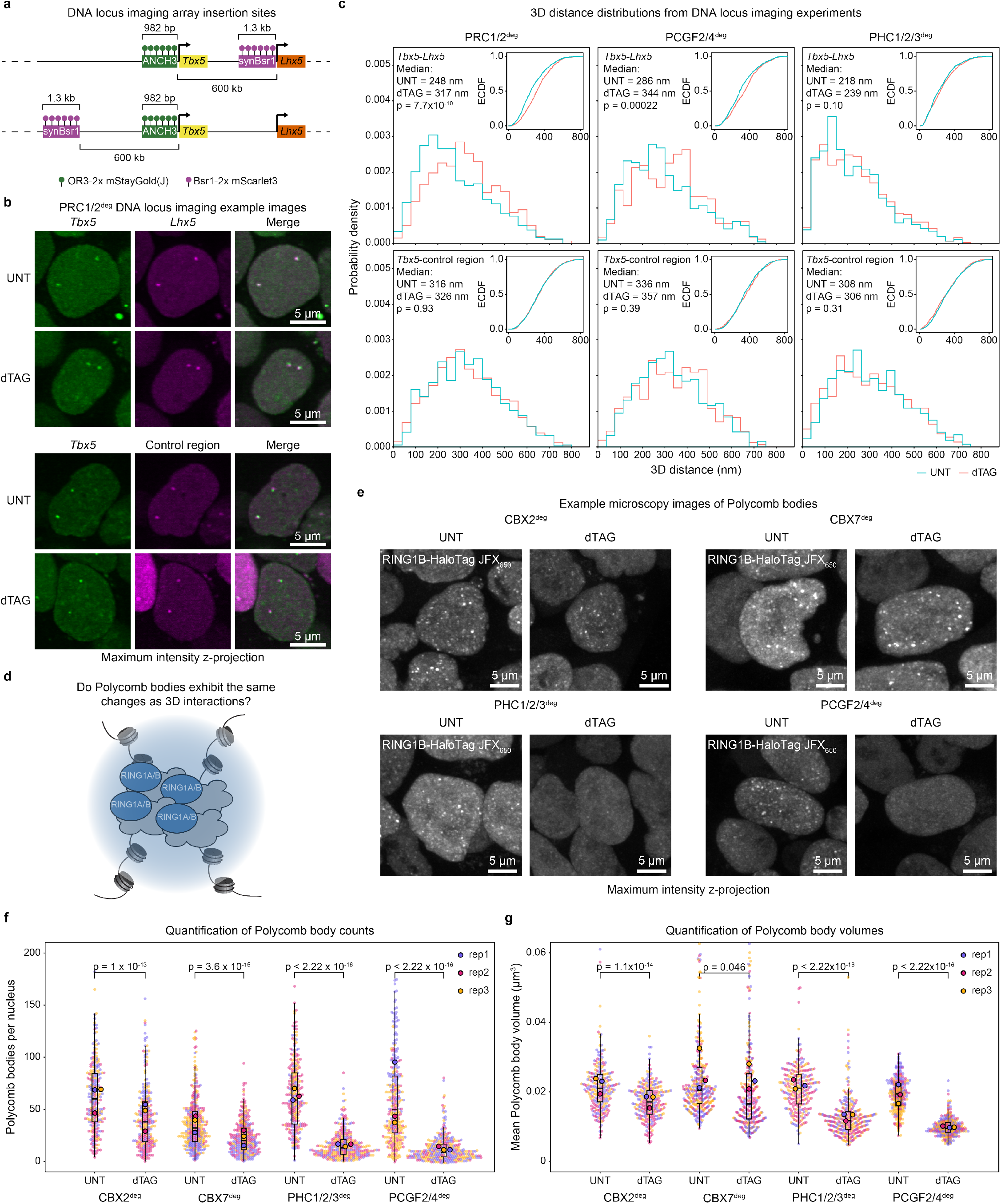
cPRC1 depletion increases the distance between Polycomb-occupied genes in live-cell imaging and disrupts Polycomb bodies. **a**, A schematic showing the design of array and binding protein insertions at the *Tbx5, Lhx5*, and negative control regions in PRC1/2^deg^ and PHC1/2/3^deg^ cell lines to allow for live-cell distance measurements between Polycomb-occupied loci using fluorescence microscopy. **b**, Example chromatic aberration-corrected maximum intensity z-projections of PRC1/2^deg^ cells with arrays inserted at *Tbx5* and *Lhx5* or at *Tbx5* and a control region, with or without dTAG-13 treatment. Two arrays are visible as foci per nucleus. **c**, Probability density plots and corresponding empirical cumulative density function (ECDF) plots (insets) for pairwise 3D distance distributions for the *Tbx5-Lhx5* and *Tbx5*-negative control loci in PRC1/2^deg^ PCGF2/4^deg^, and PHC1/2/3^deg^ in untreated (UNT) or 4 h dTAG-13 treated (dTAG) cells. Distributions represent data from three biological replicates for each condition (Number of observations per sample: PRC1/2^deg^ *Tbx5-Lhx5* UNT *n* = 583; PRC1/2^deg^ *Tbx5-Lhx5* dTAG *n* = 631; PRC1/2^deg^ *Tbx5-*control region UNT *n* = 1089; PRC1/2^deg^ *Tbx5-*control region dTAG *n* = 901; PCGF2/4^deg^ *Tbx5-Lhx5* UNT *n* = 485; PCGF2/4^deg^ *Tbx5-Lhx5* dTAG *n* = 399; PCGF2/4^deg^ *Tbx5-*control region UNT *n* = 476; PCGF2/4^deg^ *Tbx5-*control region dTAG *n* = 394; PHC1/2/3^deg^ *Tbx5-Lhx5* UNT *n* = 645; PHC1/2/3^deg^ *Tbx5-Lhx5* dTAG *n* = 704; PHC1/2/3^deg^ *Tbx5-*control region UNT *n* = 665; PHC1/2/3^deg^ *Tbx5-*control region dTAG *n* = 549). Indicated *p*-values were calculated using the Kolmogorov-Smirnov test comparing the UNT and dTAG distributions. **d**, A cartoon model of a Polycomb body overlaid on a Polycomb-mediated interaction. **e**, Example confocal microscopy maximum intensity z-projections of RING1B-HaloTag labeled with JFX_650_ in CBX2^deg^, CBX7^deg^, PHC1/2/3^deg^, and PCGF2/4^deg^ nuclei. Scale bars represent 5 µm. **f-g**, Plots showing the distribution of the number of Polycomb bodies detected per nucleus (**f**), mean Polycomb body volume in each nucleus (**g)** for CBX2^deg^, CBX7^deg^, PHC1/2/3^deg^, and PCGF2/4^deg^ nuclei. Individual values for each nucleus are colored by treatment replicate (small circles) and are overlaid with a boxplot of the distribution of the individual nuclei and the means of all nuclei in each of three biological replicates (large, outlined circles). Number of nuclei: CBX2^deg^ UNT *n* = 252; CBX2^deg^ dTAG *n* = 256; CBX7^deg^ UNT *n* = 343; CBX7^deg^ dTAG *n* = 303; PHC1/2/3^deg^ UNT *n* = 265; PHC1/2/3^deg^ dTAG *n* = 307; PCGF2/4^deg^ UNT *n* = 303; PCGF2/4^deg^ dTAG *n* = 325. For boxplots, boxes represent the interquartile range (IQR), the middle line corresponds to the median, and whiskers extend to the largest and smallest values no more than 1.5 x IQR from the box. Indicated *p*-values were calculated using the Mann-Whitney U Test on per-nucleus values.

We performed live-cell imaging of PRC1/2^deg^, PHC1/2/3^deg^, and PCGF2/4^deg^ cells with and without 4 h dTAG-13 treatment and measured the resulting distribution of 3D distances^87^ between either *Tbx5* and *Lhx5* or *Tbx5* and the negative control region. As expected, the distance remained unchanged for the negative control interaction in all degrons (Fig. 4c). By contrast, the median distance between *Tbx5* and *Lhx5* increased by 69 nm in PRC1/2^deg^ following dTAG-13 treatment. Similarly, we found that depletion of PCGF2/4 led to an increase in median distance of 58 nm. However, despite weakened interactions in RCMC in PHC1/2/3^deg^, the distance distribution between *Tbx5* and *Lhx5* was not significantly changed following dTAG-13 treatment (Fig. 4c). This difference between PHC1/2/3^deg^ and PCGF2/4^deg^ suggests that the lack of change in distance observed in PHC1/2/3^deg^ likely results from PHC proteins remaining on chromatin (Fig. 2i).

Despite similar changes in median distance between PRC1/2^deg^ (69 nm) and PCGF2/4^deg^ (58 nm) we noted that separation increased in PRC1/2^deg^ starting at lower distances (Fig. 4c). Micro-C approaches are generally thought to capture interactions within a ≲ 50 nm radius^87,89^. Although distances measured by microscopy cannot be directly compared due to localization uncertainty^90,91^, an increase in separation at longer distances (> 100 nm) will not be as strongly detectable in RCMC due to its inability to detect interactions at that distance. Notably, only PRC1/2^deg^ changes substantially at shorter distances in our super-resolution microscopy, suggesting a difference in effect between PRC1/2^deg^ and PCGF2/4^deg^. Live cell imaging of interacting Polycomb loci therefore is consistent with the weaker effect on Polycomb-mediated interactions of PHC1/2/3^deg^ and PCGF2/4^deg^ compared to PRC1/2^deg^ seen in our RCMC data.

### Polycomb bodies are more sensitive to cPRC1 depletion than Polycomb-mediated interactions

Polycomb bodies are nuclear foci of Polycomb proteins which have been previously found to overlap interacting Polycomb-occupied loci^16^, and have been broadly proposed as the microscopically visible consequence of such interactions (Fig. 4d)^58,62,63^. We reasoned that examining the effect of cPRC1 subunit depletion on Polycomb bodies would therefore provide another readout for comparing the effect of PRC1 subunit depletions on Polycomb-mediated interactions. We used an endogenous, homozygous RING1B-HaloTag^65,92^ (Fig. S8a) inserted in all of our cell lines (with the exception of PRC1^deg^ and PRC1/2^deg^) to visualize Polycomb bodies (Fig. 4e). We imaged live cells with or without 4 h of dTAG-13 treatment and segmented Polycomb bodies from deconvolved images. We then analyzed features of the Polycomb bodies in each condition (Fig. S8b), similar to a previously described approach^65^. Although there is some cell line-to-cell line variability, we typically counted 40-60 Polycomb bodies per cell, with mean volumes of around 0.02 µm^3^ (equivalent to a sphere of ∼300 nm diameter) (Fig. 4f).

Consistent with the results from RCMC, PHC1/2/3^deg^, PHC1/2/3 CBX2/7^deg^, and PCGF2/4^deg^ exhibited a much stronger reduction in the mean number of Polycomb bodies per nucleus than CBX2^deg^ or CBX7^deg^ (Fig. 4f and Fig. S8c). However, CBX2^deg^ and CBX7^deg^ still exhibited small effects on Polycomb body counts. PHC1/2/3, PHC1/2/3 CBX2/7, and PCGF2/4 depletion also caused a large decrease in Polycomb body volume, and a slight reduction in the enrichment of PRC1 within Polycomb bodies, compared to depletion of CBX2 or CBX7 (Fig. 4g and Fig. S8d,e). Notably, we also observed that total RING1B signal was reduced by 10% following PHC1/2/3 depletion and by 25% following PCGF2/4 depletion (Fig. S8f). This suggests there is a general disruption of cPRC1 complex stability not only in the absence of PCGF2/4, as previously reported^70^, but also when PHC proteins are depleted. This observation is consistent with a recent structural study showing an integral role for PHC proteins in cPRC1 complexes^93^. Together, we observed similar relative effects of cPRC1 subunit depletion on Polycomb bodies as in RCMC, yet at an apparent higher magnitude. This suggests that Polycomb bodies are more sensitive to cPRC1 levels than Polycomb-mediated interactions observed in RCMC.

We also note that the scale of Polycomb bodies (∼300 nm) closely matches the length scale of altered distances between *Tbx5* and *Lhx5* in our super-resolution imaging (Fig. 4c). If Polycomb bodies do represent Polycomb-occupied loci in proximity, this may provide a possible explanation for increased separation at longer distances, while shorter range interactions are maintained by alternative mechanisms.

### cPRC1-independent Polycomb-mediated interactions do not depend on a single specific complex

To date, Polycomb-mediated interactions have generally been described as the result of protein-protein interactions depending on cPRC1 subunits^11,16,58,62,63^. On this basis, we wondered whether the remaining interaction strength might arise from previously undescribed interaction capability of other Polycomb complexes. Therefore, we focused on the Polycomb complexes that are intact in the absence of cPRC1 but disrupted in the PRC1/2^deg^ cell line. There are three classes of vPRC1 complexes which do not contain PCGF2/4: PRC1.1 (containing PCGF1), PRC1.3/1.5 (containing PCGF3/5), and PRC1.6 (containing PCGF6)^27,29,30^. Of these, PRC1.3/1.5 generally lack localized or stable chromatin binding^65,70,94^, and do not overlap with sites which form Polycomb-mediated interactions. In addition to vPRC1, PRC2 complexes are also intact in cPRC1 depletion conditions. While we have shown that PRC1 depletion alone is equivalent to PRC1/2 depletion (Fig. 1), we reasoned that this effect could be due to the loss of PRC2 chromatin occupancy that follows depletion of PRC1 and H2AK119ub1^44,70,71,82^. We also determined that depletion of SUZ12 (PRC2^deg^) also had an impact on Polycomb-mediated interactions (Fig. S5), which could be direct or indirect. We therefore generated three additional triple degron cell lines based on our PCGF2/4^deg^ line, tagging SUZ12 (PCGF2/4 SUZ12^deg^), PCGF1 (PCGF1/2/4^deg^), and PCGF6 (PCGF2/4/6^deg^) (Fig. 5a and Fig. S9a,b,c,d).

**Figure 5.**
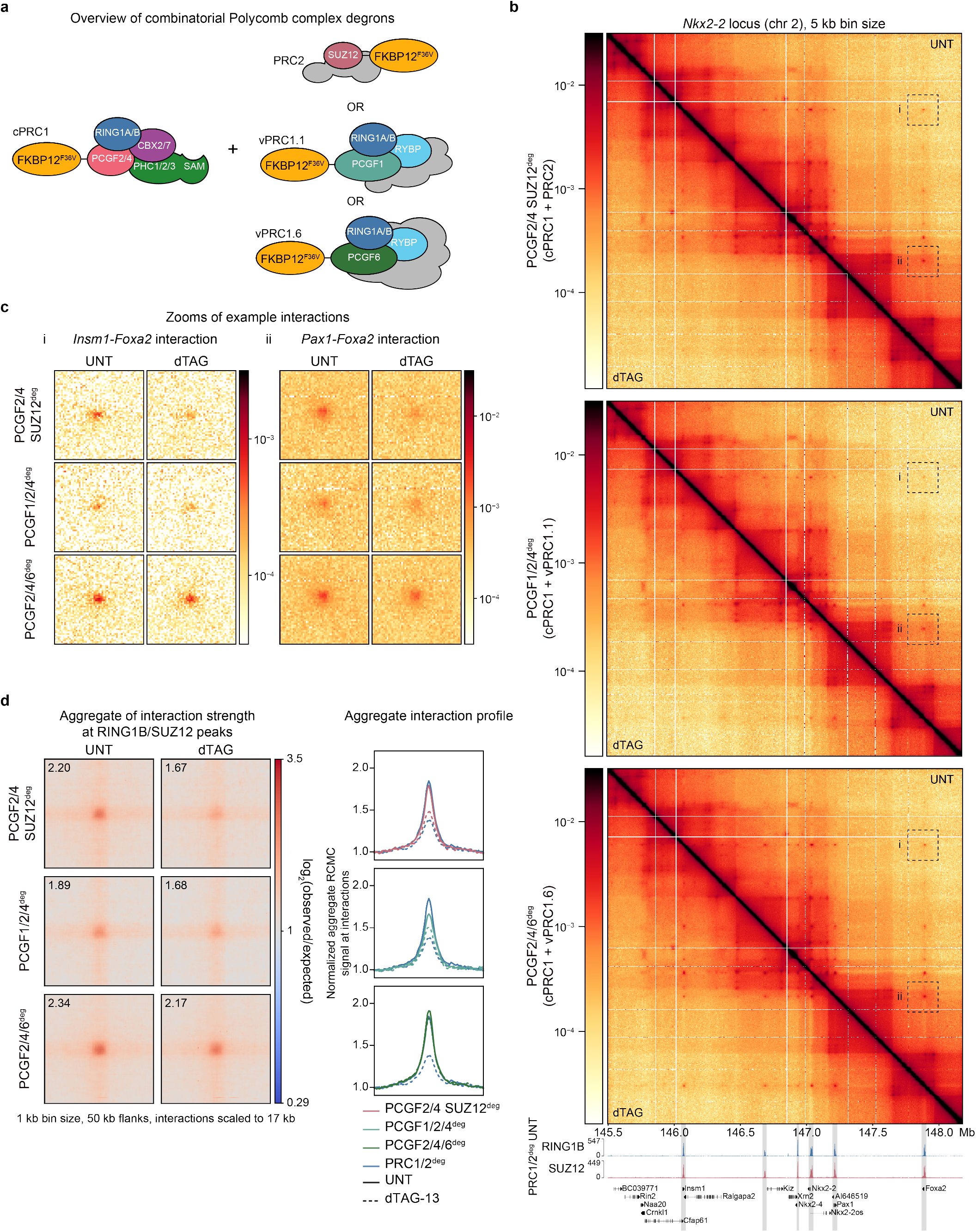
Disruption of individual PRC1 and PRC2 complexes is insufficient to fully disrupt Polycomb-mediated interactions. **a**, A schematic illustrating combinatorial degrons of PCGF2/4 with SUZ12, PCGF1, or PCGF6. **b**, RCMC log_10_-scaled contact maps for PCGF2/4 SUZ12^deg^, PCGF1/2/4^deg^, and PCGF2/4/6^deg^ showing changes in contact frequency (upper) with corresponding RING1B and SUZ12 ChIP-seq in PRC1/2^deg^ UNT illustrating the locations of Polycomb-occupied regions. Data shown are merges of *n* = 2 (RCMC) or *n* = 3 (ChIP-seq) biological replicates. **c**, Zooms of interactions from RCMC contact map regions labeled i and ii in **b** in untreated and dTAG-13 treatment, labeled according to the genes at the interacting loci. Contact map zooms are plotted at 2 kb bin size. **d**, Left panel: aggregate pileups of interactions (log_2_(observed/expected)) between RING1B/SUZ12 co-occupied sites in degron lines in **b** with or without 4 h dTAG-13 treatment. Sites are scaled to 17 kb with 50 kb flanks, with data plotted at 1 kb bin size. Values in the upper left corner of each plot indicate the mean value of the central 3×3 pixels of the pileup. Right panel: a 1D average plot across the 21 middle diagonals of the aggregate pileups in the left panel and **Fig. 1f** for PRC1/2^deg^, normalized to the mean of the 10 bins on the left- and right-most sides.

As expected for a SUZ12 depletion, the PCGF2/4 SUZ12^deg^ cell line resulted in a decrease in H3K27me3 not seen for PCGF2/4^deg^ (Fig. S9e). Depletion of PCGF1/2/4 resulted in a slightly higher number of derepressed genes than PCGF2/4 alone or PCGF2/4/6, consistent with previous studies showing PCGF1 has a larger impact on transcription^70,94,95^ (Fig. S9f). As in the parental PCGF2/4^deg^ cell line, each of these lines exhibited a near-complete disruption of Polycomb bodies (Fig. S9g,h,i) and a reduction in total RING1B (Fig. S9j). However, none of the three degrons reduced the strength of Polycomb-mediated interactions in RCMC to the same degree as in PRC1/2^deg^ (Fig. 5b,c,d). PCGF2/4 SUZ12^deg^ and PCGF1/2/4^deg^ showed similar interaction strength after dTAG-13 treatment to the parental PCGF2/4^deg^ cell line, suggesting no additional effect of the added depletion of SUZ12 or PCGF1. In the PCGF2/4/6^deg^ cell line, the reduction in interaction strength was even weaker than in the PCGF2/4^deg^ line. This is possibly consistent with the reduced number of derepressed genes in PCGF2/4/6^deg^ compared to PCGF2/4^deg^, suggesting a compensatory effect of PCGF6 depletion.

Altogether, no individual Polycomb complex with specific genomic targeting can account for the residual Polycomb-mediated interaction strength following cPRC1 depletion. This suggests that the residual interaction strength may be mediated by affinity, such as between Polycomb proteins generally or the chromatin state at Polycomb-occupied genes (Fig. 6).

**Figure 6.**
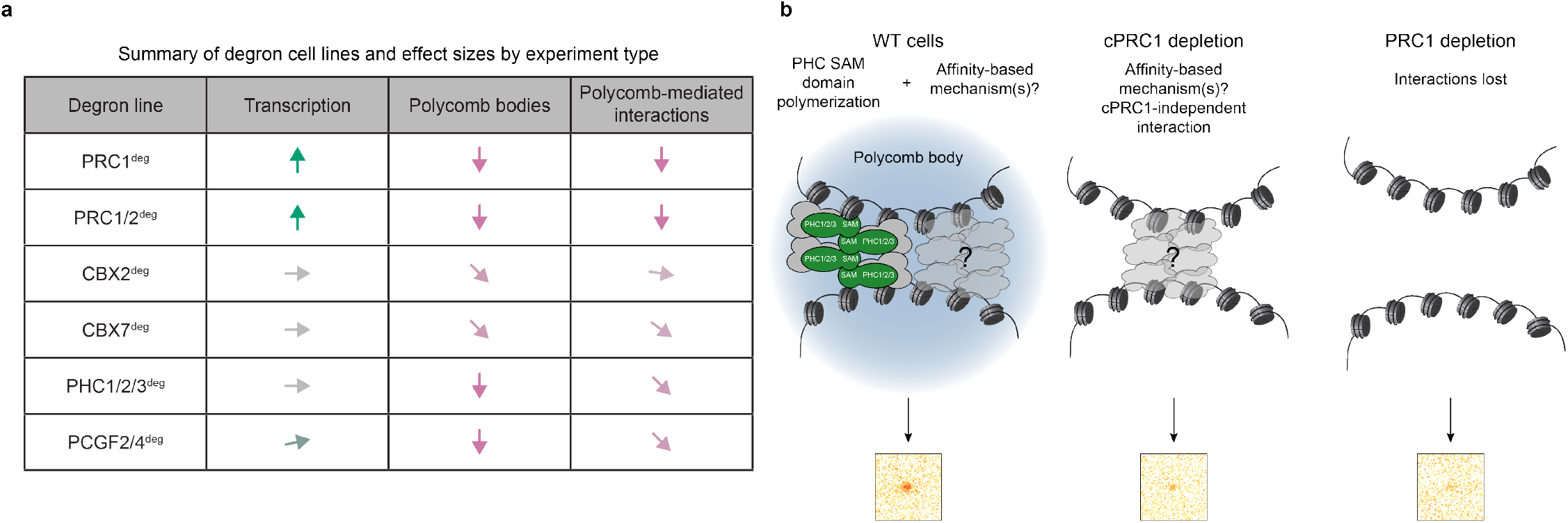
Depletion of cPRC1 is sufficient to disrupt Polycomb bodies but Polycomb-mediated interactions can persist in its absence. **a**, A table summarizing experimental results for the six major degron cell lines used in this study in terms of the effect on transcription, Polycomb bodies, and Polycomb-mediated interactions. Arrow orientation and color correspond qualitatively to effect size. **b**, A proposed model for Polycomb-mediated interactions in mESCs, whereby interactions are formed in WT cells by a combination of PHC proteins in cPRC1 complexes and a potentially more general, affinity-mediated interaction, leading to Polycomb body formation and strong Polycomb-mediated interactions by RCMC. In this model, depletion of PHC proteins or cPRC1 leads to a loss of cPRC1-mediated interaction strength, but a retention of interactions due to this residual unknown mechanism. Finally, depletion of Polycomb in its entirety leads to a loss of interactions.

## DISCUSSION

In this work, we have carried out a comprehensive dissection of the protein factors driving Polycomb-mediated 3D chromatin interactions and Polycomb body formation in mESCs. We find that depletion of PRC1 alone is sufficient to disrupt interaction formation to the same extent as PRC1 and PRC2 depletion (Fig. 1). Using depletion cell lines for specific cPRC1 subunits, we find that CBX2 makes only a minimal contribution to Polycomb-mediated interactions and a modest contribution to Polycomb bodies, while PHC proteins drive a large proportion of these interactions (Fig. 2 and Fig. 4). By extending our depletions to encompass the entirety of cPRC1, we show that a small proportion of Polycomb-mediated interaction strength is not explained by cPRC1 complexes (Fig. 3). We further deplete three other Polycomb complexes in combination with cPRC1, but find that none of these is sufficient to explain Polycomb-mediated interactions in the absence of cPRC1 (Fig. 5). Together, these results provide a detailed view of the roles of cPRC1 subunits, their contributions to PRC1 chromatin occupancy, Polycomb-mediated interactions, and Polycomb bodies, and reveal an unexpected finding that Polycomb-mediated interactions can exist independently of known cPRC1 complexes in mESCs (Fig. 6).

Two major models for cPRC1 contribution to Polycomb interactions have previously been proposed, one driven by the self-interaction of the CaPS domain in CBX proteins^58^, and the other by PHC SAM domain polymerization^11,16^. We determine that CBX2 makes a negligible contribution to Polycomb-mediated interactions and a smaller contribution to Polycomb bodies than PHC proteins in mESCs. This is not entirely unexpected: previous studies have found that PHC proteins contribute strongly to Polycomb-mediated interactions and Polycomb bodies in mESCs^11,16^, while CBX2 is very lowly expressed^58^. Nonetheless, recent work has focused on CBX2 and its condensate-forming activity as a potential regulator of interactions in mESCs, despite its low expression^58^. In other cell types, where expression of CBX7 decreases and other, CaPS domain-containing CBX proteins are more highly expressed, we expect that CBX proteins may play a larger role in Polycomb mediated interactions^29,59,68^. Examination of Polycomb complex composition and 3D chromatin interactions in more differentiated cell types will be required to uncover how these contributions change and the consequences for Polycomb system function.

The prevailing view is that cPRC1 is solely responsible for Polycomb-mediated interactions^11,16,58,62,63^, and evidence suggests that cPRC1 binding is sufficient to form interactions^57^. However, we unexpectedly find that cPRC1 depletion cannot recapitulate the effect of PRC1 depletion on Polycomb-mediated interactions. We find it likely that this finding is possible due to a combination of the high sensitivity of RCMC and our comparative measurements of interaction strength. Despite our efforts to perturb this residual interaction by perturbation of remaining Polycomb complexes (PRC1.1, PRC1.6, PRC2), we are unable to pinpoint from where it arises. This leads us to speculate that this interaction strength results from one of two molecular mechanisms. First, it is possible that unknown subunits or protein-protein interactions of known subunits could mediate these residual interactions. We think this is a less likely possibility, as this would likely require at least one Polycomb complex, and would therefore have been disrupted in one of the PCGF2/4 SUZ12^deg^, PCGF1/2/4^deg^, or PCGF2/4/6^deg^ conditions. A notable exception is RYBP, which is present across all vPRC1 complexes and allows PRC1 to bridge adjacent nucleosomes^96,97^. However, RYBP has an unusually short chromatin residence time (∼15 s) compared to other PRC1 subunits, and appears less stably associated with the complex^65^, making it an unlikely structural factor. Second, analogously to enhancer-promoter affinity-mediated interactions^4,80,98,99^, we propose that less specific, affinity-based mechanisms could drive cPRC1-independent interaction strength. PRC1 subunits could have general affinity for each other, requiring total PRC1 depletion to disrupt interactions. Alternatively, H2AK119ub1, or other features of Polycomb-repressed promoters could drive affinity between regions of similarly marked chromatin. Depletion of PRC1 leads to rapid loss of H2AK119ub1 and derepression of many Polycomb-occupied genes, changing promoter state and potentially disrupting such affinity.

Any mechanism for cPRC1-independent interactions requires the involvement of vPRC1 complexes either directly or for H2AK119ub1 deposition and transcriptional repression. Invoking the involvement of vPRC1 complexes is appealing as it would help resolve an apparent contradiction of cPRC1-driven interactions: that cPRC1 has a minimal effect on transcription^70^, while perturbations which weaken or strengthen Polycomb-mediated interactions lead to changes in gene expression^21,69^. If vPRC1, or the chromatin state it generates, also plays a role in Polycomb-mediated interactions, this would explain the observed link between Polycomb-mediated interactions and transcription. Extensive further work and careful study design will be needed to unpick the interdependencies of these features to determine a mechanism.

Alongside our study of 3D genome interactions, we also determine the effect that depletion of various Polycomb proteins has on Polycomb bodies. In contrast to the changes we observe in RCMC, Polycomb bodies exhibit near-complete loss following disruption of cPRC1. This observation is consistent with more than one possible explanation. This might suggest that Polycomb bodies do not represent sites of interaction between highly Polycomb-occupied sites, and are instead clusters of self-interacting cPRC1. This explanation seems unlikely given existing evidence that strong Polycomb target genes are often localized inside Polycomb bodies^16,63^. Furthermore, other Polycomb subunits, such as PCGF1 and PCGF6, also localize to Polycomb bodies^65^. Another possibility is that Polycomb bodies represent Polycomb-mediated interactions, but depend more on cPRC1 and cease to be visible without cPRC1. Regardless of the explanation, Polycomb bodies can be perturbed without impacting transcription in mESCs, indicating either a redundant role or that they are a passive consequence of cPRC1 self-interaction, unrelated to transcriptional repression.

Recent work reported that CBX7 chromatin binding depends on PRC2^52,53^. In line with this, we find that PRC2^deg^ weakens Polycomb-mediated interactions similarly to cPRC1 depletion, despite only partial reduction in H3K27me3 at 4 h. We also observe a reciprocal effect, where CBX7 depletion reduces SUZ12 chromatin occupancy. This raises the possibility that CBX7 can also support PRC2 chromatin binding, providing a potential function for cPRC1 beyond forming interactions and compacting chromatin.

Altogether, our comparison of multiple Polycomb subunit degrons reveals that, while PHC is the main driver of Polycomb-mediated interactions within cPRC1, interactions can unexpectedly be maintained in the absence of cPRC1. This provides an intriguing potential link between Polycomb-mediated interactions and gene repression which has not been explored previously, the mechanism of which remains to be elucidated.

## Acknowledgements

We thank Rob Klose for support and input in setting up the project. We also thank Rob Klose and Emilia Dimitrova for sharing PCGF2 antibodies for ChIP-seq experiments. We are further grateful to Emilia Dimitrova and Amy Hughes for guidance and support on TT_chem_-seq. We thank Leonid Mirny for discussions of data and conclusions. We thank Sarah Nemsick, James Jusuf, Clarice Hong, Sushanta Mahanta, Domenic Narducci, and Emma Magee for critical reading and feedback on the manuscript. We further thank all members of the Hansen lab for constructive input and support throughout the work. We thank the MIT Koch

Institute’s Robert A. Swanson (1969) Biotechnology Center for technical support, specifically the Integrated Genomics and Bioinformatics Core, MIT BioMicroCenter, and Flow Cytometry Core. We also thank the Walk-Up Sequencing services of the Broad Institute of MIT and Harvard. We thank the High-Content Imaging Facility team at the Center for Development of Therapeutics at the Broad Institute for maintaining the Opera Phenix High-Content Screening System.

## Author Contributions

M.K.H. designed the project with input from A.S.H. and S.K. N.P.B. generated and validated the parental inducible *Ngn2* mESC line and the PRC1^deg^ cell line. M.K.H. generated the RING1B-HaloTag line and generated and validated all other degron cell lines. S.K. generated cell lines including locus labels for single-locus imaging. M.K.H. performed and analyzed all genomics experiments. M.K.H. performed imaging of Polycomb bodies, S.K. developed the Polycomb body image analysis with assistance from J.T. S.K. performed Polycomb body image processing and quality control, and M.K.H. analyzed pipeline outputs. S.K. performed locus imaging experiments, J.T. modified Fyrtarn for two color processing, S.K. added minor modifications to process single timepoints, and S.K. performed the analysis. M.K.H performed and analyzed all other experiments. M.K.H. drafted the manuscript and generated the figures with input from all authors. S.K. contributed to microscopy figure panels. A.S.H. supervised the research. All authors contributed to editing the manuscript and figures.

## Data Availability

All genomics data generated in this study are publicly available on the NCBI Gene Expression Omnibus in SuperSeries GSE341617. Previously published ATAC-seq data is available at GSE98403, and previously published Hi-C data is available at ArrayExpress E-MTAB-7816. Previously published ChIP-seq datasets are available as follows: H3K4me3 GSE199801, PCGF1 GSE132753, PCGF6 GSE132753, P300 GSE90893, BRD4 GSE87037, BRG1 GSE90893, CBX3 GSE44242, H2AZ GSE34483, H3K4me1 GSE90893, H3K9me2 GSE54412, H3K9me3 GSE90893, H3K27ac GSE90893, MED1 GSE22562, MED12 GSE22562, CTCF GSE90994, RAD21 GSE90994, SMC1A GSE123636. Outputs from 3D distance measurements from microscopy and Polycomb body segmentation are available at https://doi.org/10.5281/zenodo.23224995.

## Code Availability

All new custom code and scripts described in this paper are available at https://github.com/ahansenlab/Polycomb_RCMC_analysis_code for code used in analysis, and at https://github.com/ahansenlab/Polycomb_body_image_processing for code used for image processing of Polycomb body microscopy.

## Materials and Methods

Materials and methods can be found in the Supplementary Information.

