## Supplementary Information for "Dissection of Polycomb complexes reveals canonical PRC1-independent 3D chromatin interactions"

### Supplementary information for
### Dissection of Polycomb complexes reveals canonical PRC1-independent 3D chromatin interactions

Miles K. Huseyin[1,2,3,4,*], Sumin Kim[1,2,3,4], Jack Toppen[1,2,3,4], Neil P. Blackledge[5], Anders S. Hansen[1,2,3,4,*]

[1]Department[1] Department of Biological Engineering, Massachusetts Institute of Technology; Cambridge, MA 02139, USA; [2] Gene Regulation Observatory, Broad Institute of MIT and Harvard; Cambridge, MA 02142, USA; [3] The Novo Nordisk Foundation Center for Genomic Mechanisms of Disease, Broad Institute of MIT and Harvard; Cambridge, MA 02142, USA; [4] Koch Institute for Integrative Cancer Research; Cambridge, MA, 02139, USA[5]; Department of Biochemistry, University of Oxford; Oxford OX1 3QU, UK

#### Contents

- Supplementary Figures & Figure Legends
- Materials and Methods
- Supplementary Tables
- References for Supplementary Information

### SUPPLEMENTARY FIGURES & FIGURE LEGENDS

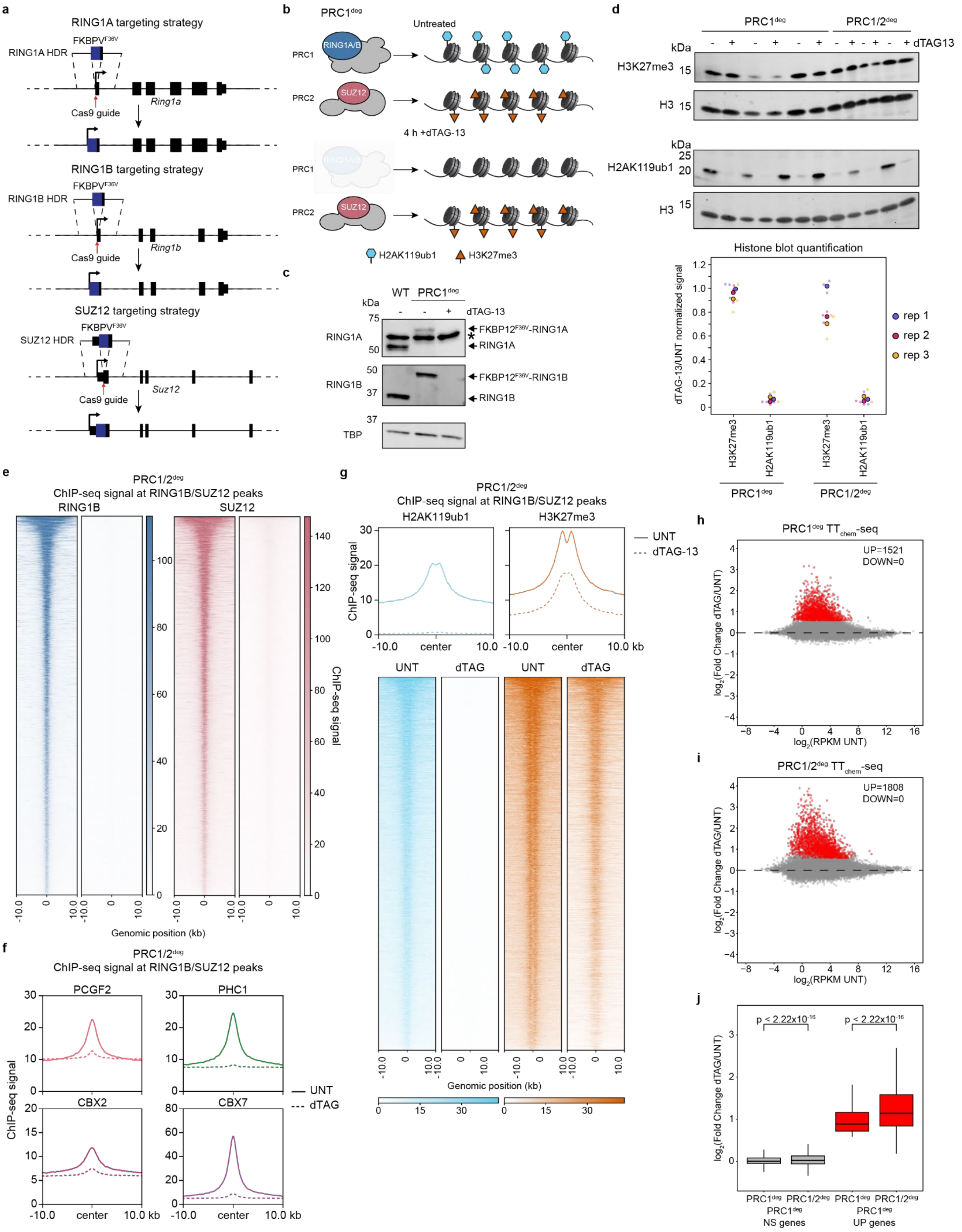

**Supplementary Figure 1. Generation of PRC1 and PRC1/2 degron cell lines. a,** A schematic illustrating the genome editing strategy for inserting the FKBPV$^{F36V}$ degron tag into the *Ring1a*, *Ring1b*, and *Suz12* genes. Narrow boxes indicate untranslated sequences, while wider boxes are translated exons. **b,** A schematic illustrating the Polycomb complexes and histone modifications in untreated and 4 h dTAG-13 treated PRC1$^{deg}$ (RING1A/B degron) cells. **c,** Western blot validation of RING1A and RING1B depletion following 4 h dTAG-13 treatment in PRC1$^{deg}$ cell lines. TBP is included as a loading control. Asterisk indicates a non-specific band. **d,** Upper panel: representative western blots from one technical replicate of Polycomb-mediated histone modifications H2AK119ub1 and H3K27me3 from histone extracts showing three replicates of 4 h treatment with dTAG-13 for each of PRC1$^{deg}$ and PRC1/2$^{deg}$. Histone H3 is included as a loading control. All three technical replicates of western blotting are provided as uncropped blots. Lower panel: quantification of histone western blots for PRC1deg and PRC1/2 deg treatments, normalized for corresponding histone H3 signal. Small circles represent technical replicates of western blotting, while large circles represent the means of the technical replicates for each treatment replicate. All plotted data are colored according to their treatment replicate. **e,** Heatmaps showing loss of RING1B and SUZ12 ChIP-seq signal at RING1B/SUZ12 co-occupied peaks following 4 h dTAG-13 treatment in PRC1/2$^{deg}$. Data represent the merge of *n* = 3 biological replicates per condition. **f,** Metaplots showing loss of ChIP-seq signal for cPRC1 subunits PCGF2, PHC1, CBX2, and CBX7 at RING1B/SUZ12 co-occupied peaks following 4 h dTAG-13 treatment in PRC1/2$^{deg}$. Data represent the merge of *n* = 3 biological replicates per condition. **g,** Metaplots and heatmaps showing loss and reduction of H2AK119ub1 and H3K27me3 at RING1B/SUZ12 co-occupied peaks following 4 h dTAG-13 treatment in PRC1/2$^{deg}$. Data represent a single biological replicate for confirmation. **h-i,** MA plots of $\log_2$(fold change) in nascent gene expression measured by $TT_{chem}$-seq from UNT to dTAG in PRC1$^{deg}$ (**h**) and PRC1/2$^{deg}$ (**i**). Numbers of significantly differentially up- (UP) or down-changed (DOWN) genes are shown within each plot, and genes which are significantly changed in expression ($\log_2$(fold change) > 1.5 and $p_{adj} < 0.05$) are marked with red dots while unchanged genes are shown as grey dots. Data are derived from *n* = 3 biological replicates. **j,** Boxplots comparing $\log_2$(fold change) in gene expression between dTAG and UNT for genes which were non-significantly changed in PRC1$^{deg}$ (PRC1$^{deg}$ NS, left, grey) or significantly increased (PRC1$^{deg}$, right, red), for both PRC1$^{deg}$ and PRC1/2$^{deg}$ cell lines. Indicated *p*-values were calculated using the paired samples *t* test. Boxes represent the interquartile range (IQR), the middle line corresponds to the median, and whiskers extend to the largest and smallest values no more than 1.5 x IQR from the box.

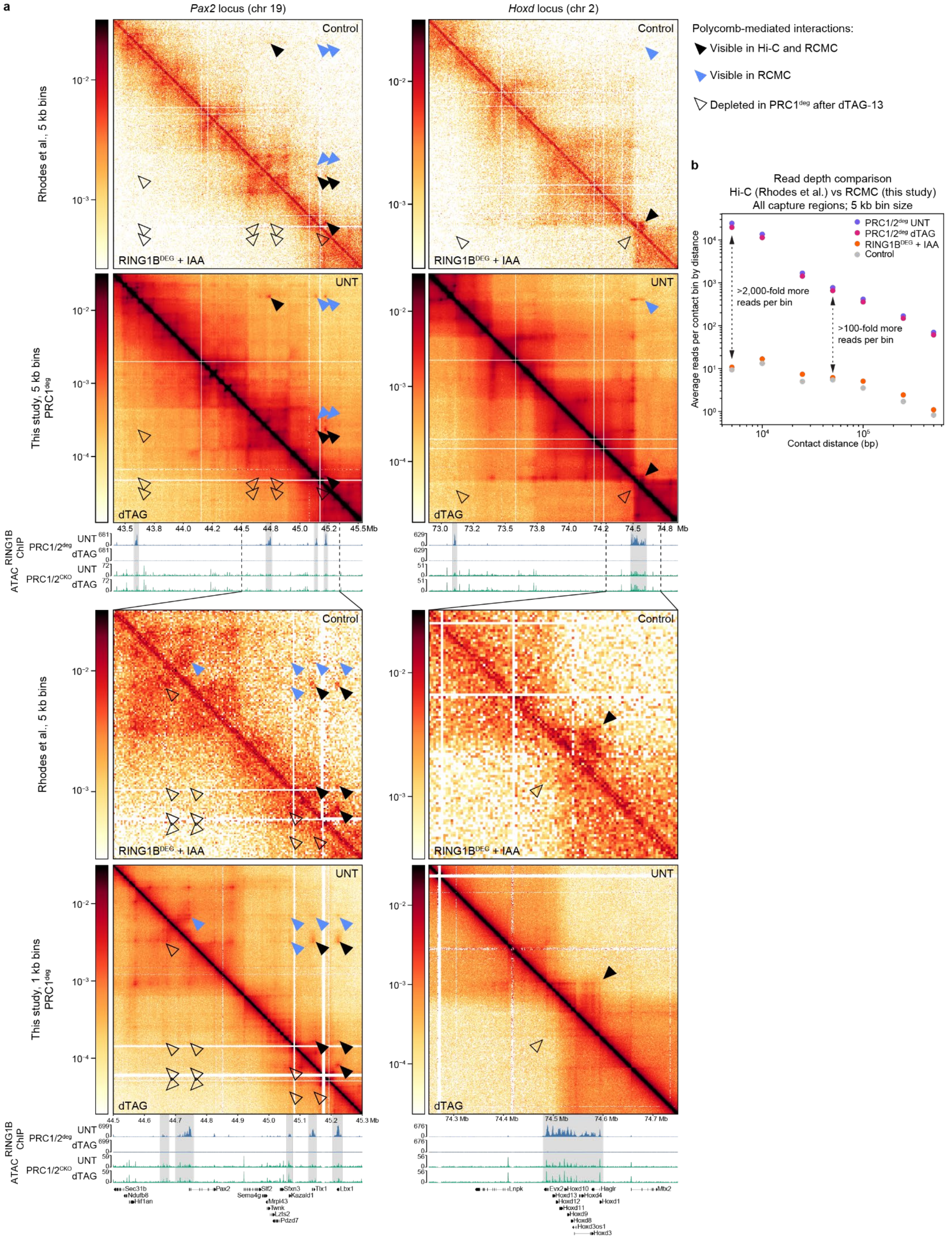
a
Pax2 locus (chr 19)
Hoxd locus (chr 2)
Polycomb-mediated interactions:
Visible in Hi-C and RCMC
Visible in RCMC
Depleted in PRC1deg after dTAG-13
Rhodes et al., 5 kb bins
This study, 5 kb bins PRC1deg
This study, 1 kb bins PRC1deg
Control
RING1BDEG + IAA
UNT
dTAG
RING1B ChIP
ATAC
PRC1/2deg
PRC1/2CKO
Contact distance (bp)
b
Read depth comparison
Hi-C (Rhodes et al.) vs RCMC (this study)
All capture regions; 5 kb bin size
Average reads per contact bin by distance
PRC1/2deg UNT
PRC1/2deg dTAG
RING1BDEG + IAA
Control
>2,000-fold more reads per bin
>100-fold more reads per bin
Sec31b
Ndufb8
Hif1an
Pax2
Sema4g
Slf2
Mrpl43
Twnk
Lzts2
Pdzd7
Sfxn3
Kazald1
Tlx1
Lbx1
Lnpk
Evx2
Hoxd13
Hoxd12
Hoxd11
Hoxd10
Hoxd9
Hoxd8
Hoxd4
Hoxd3os1
Hoxd3
Hagl
Hoxd1
Mtx2

**Supplementary Figure 2. Comparison of PRC1/2$^{deg}$ RCMC to previously published RING1B$^{deg}$ Hi-C. a,** Contact maps: Hi-C (ArrayExpress E-MTAB-7816[1]) and RCMC $log_{10}$-scaled contact maps comparing Control (Tir1 + IAA) and RING1B$^{deg}$ + IAA, or PRC1/2$^{deg}$ untreated (UNT) and 4 h dTAG-13 treated (dTAG). Genomics tracks: RING1B ChIP-seq tracks in PRC1/2$^{deg}$ cells and ATAC-seq tracks from PRC1/2$^{CKO}$ (*Ring1a*$^{-/-}$ *Ring1b*$^{fl/fl}$ *Eed*$^{-/-}$) cells (GSE98403[2]), showing a lack of change in accessibility despite long term Polycomb depletion. Regions shown in the bottom half of the figure panel are zooms of regions within the upper half, denoted by dashed and solid lines. Data shown from this study are merges of $n$ = 2 biological replicates. **b,** A comparison of reads per bin at different distances within all capture regions in PRC1/2$^{deg}$ RCMC (this study) and Hi-C data (ArrayExpress E-MTAB-7816[1]). Approximate fold differences at two contact distances are shown.

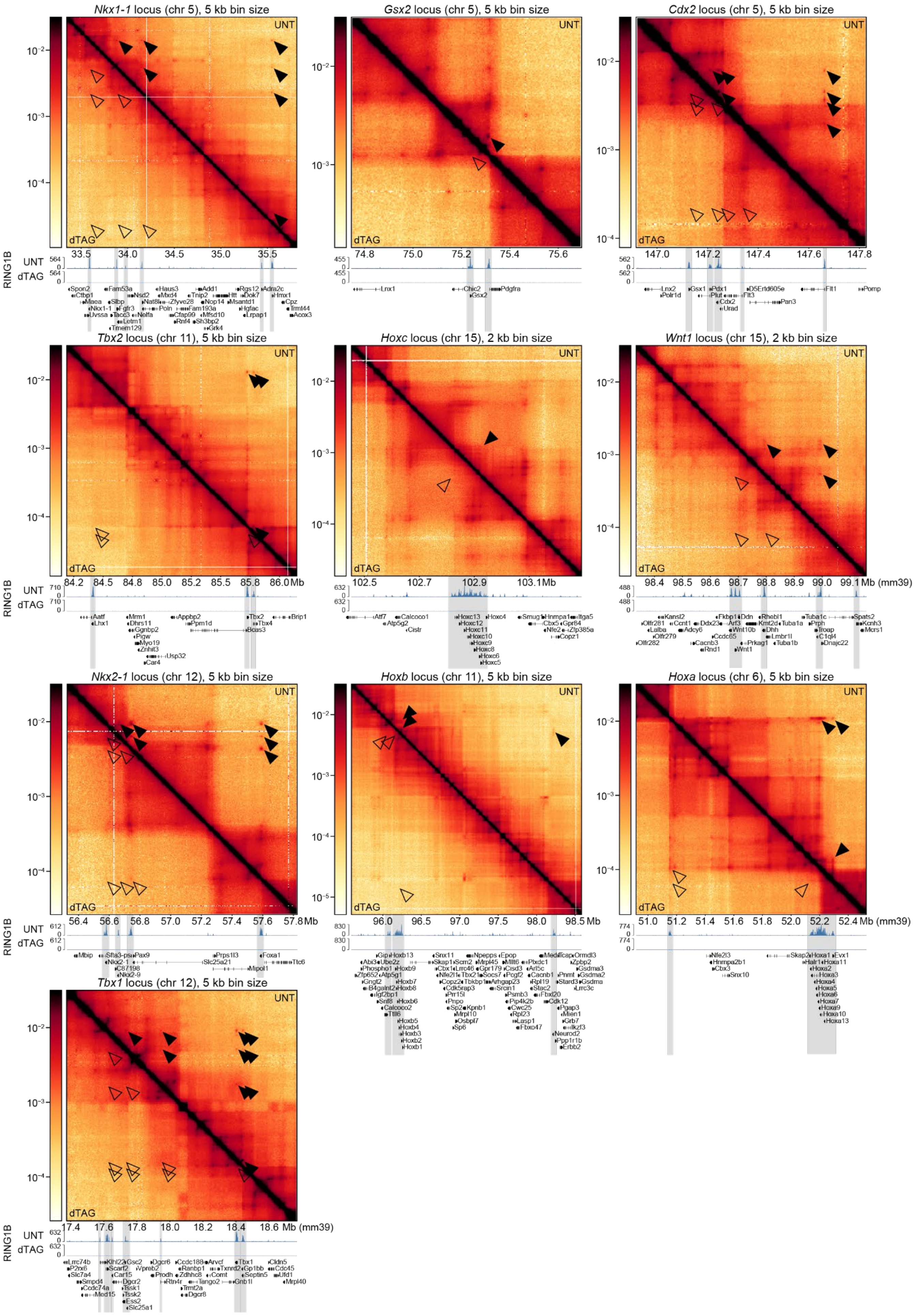

Nkx1-1 locus (chr 5), 5 kb bin size
Gsx2 locus (chr 5), 5 kb bin size
Cdx2 locus (chr 5), 5 kb bin size
Tbx2 locus (chr 11), 5 kb bin size
Hoxc locus (chr 15), 2 kb bin size
Wnt1 locus (chr 15), 2 kb bin size
Nkx2-1 locus (chr 12), 5 kb bin size
Hoxb locus (chr 11), 5 kb bin size
Hoxa locus (chr 6), 5 kb bin size
Tbx1 locus (chr 12), 5 kb bin size
UNT
dTAG
RING1B
Mb (mm39)

**Supplementary Figure 3. RCMC capture regions containing Polycomb-mediated interactions for PRC1/2$^{deg}$.** RCMC $log_{10}$-scaled contact maps for PRC1/2$^{deg}$ at all capture regions not shown elsewhere with changes in contact frequency between untreated and 4 h dTAG-13 treated cells with corresponding RING1B ChIP-seq in PRC1/2$^{deg}$ illustrating the locations of Polycomb-occupied regions. Filled arrowheads represent Polycomb-mediated interactions visible in untreated cells, while unfilled arrowheads indicate the same positions following PRC1/2 depletion. Data shown are merges of $n = 2$ (RCMC) or $n = 3$ (ChIP-seq) biological replicates.

**a**

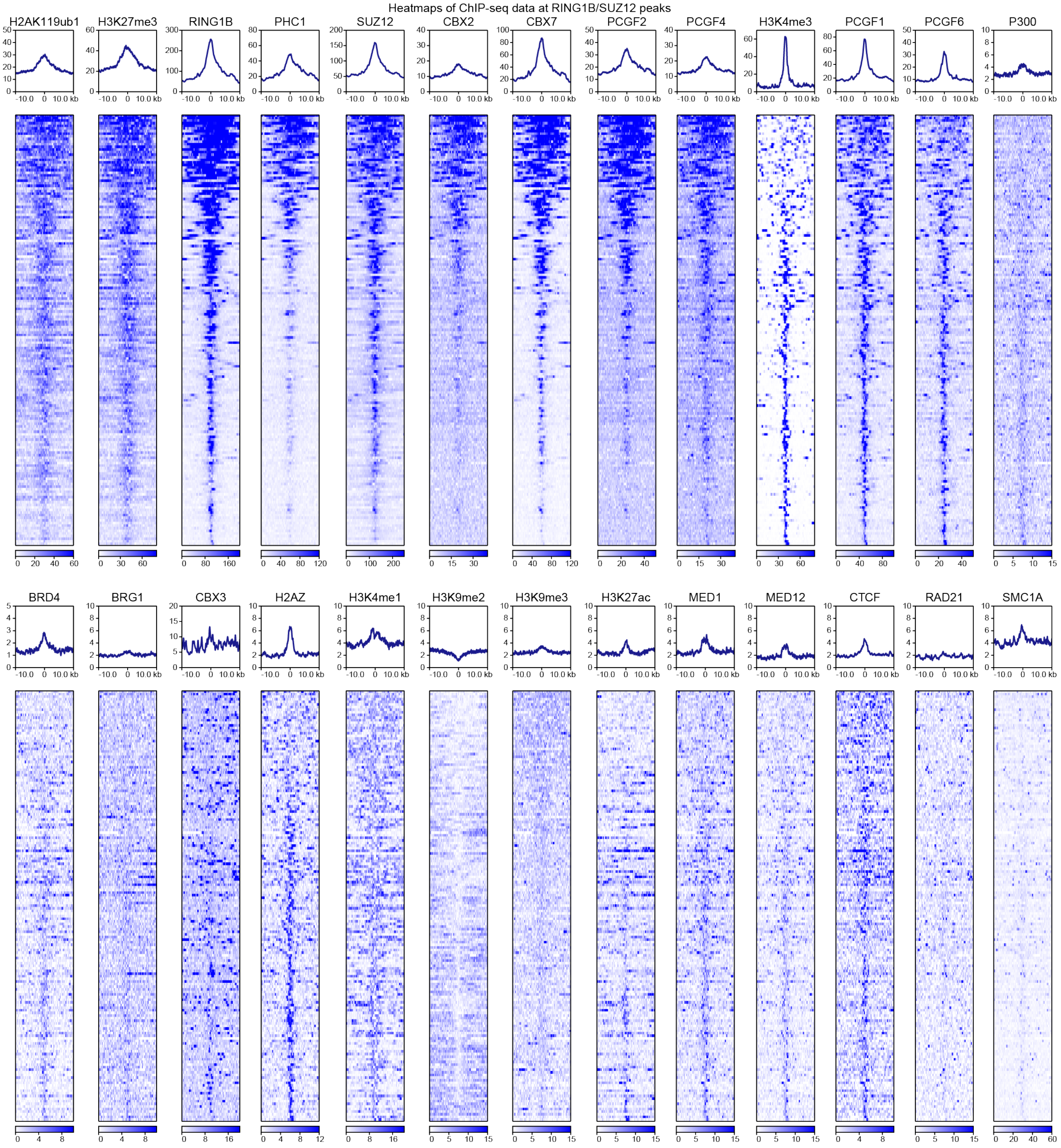


**b**

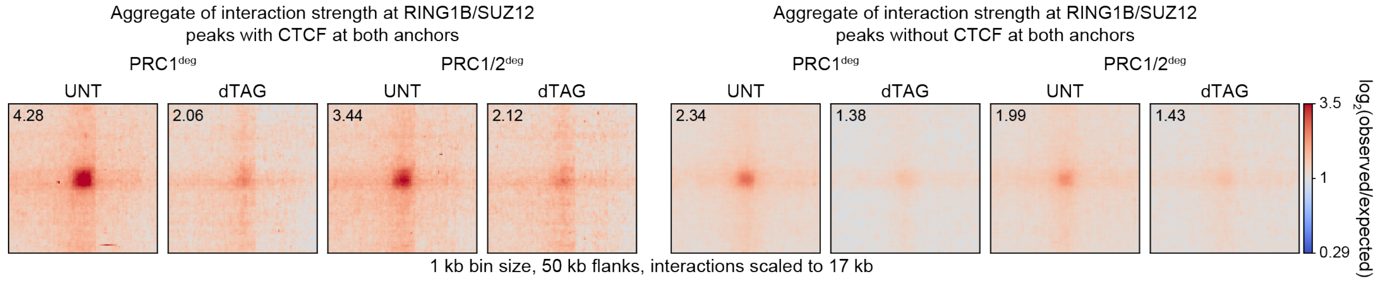


**Supplementary Figure 4. Sites of Polycomb-mediated interactions do not overlap extensively with other chromatin features. a,** Heatmaps and metaplots of assorted ChIP-seq data from this study and others centered on RING1B/SUZ12 peaks used to define interaction sites (*n* = 165). Datasets from H2AK119ub1 through PCGF2 were generated from PRC1/2$^{deg}$ cells in this work. PCGF4 ChIP-seq was generated from PCGF2/4$^{deg}$ cells. Remaining datasets are publicly available as follows: H3K4me3 GSE199801, PCGF1 GSE132753, PCGF6 GSE132753, P300 GSE90893, BRD4 GSE87037, BRG1 GSE90893, CBX3 GSE44242, H2AZ GSE34483, H3K4me1 GSE90893, H3K9me2 GSE54412, H3K9me3 GSE90893, H3K27ac GSE90893, MED1 GSE22562, MED12 GSE22562, CTCF GSE90994, RAD21 GSE90994, SMC1A GSE123636. **b,** Aggregate pileups of interactions ($log_2$(observed/expected)) between RING1B/SUZ12 co-occupied sites in $PRC1^{deg}$ and $PRC1/2^{deg}$ which overlap CTCF/SMC1 co-occupied sites at both ends (left, *n* = 100) or which do not (right, *n* = 417). Sites are scaled to 17 kb with 50 kb flanks, with data plotted at 1 kb bin size. Values in the upper left corner of each plot indicate the mean value of the central 3x3 pixels of the pileup.

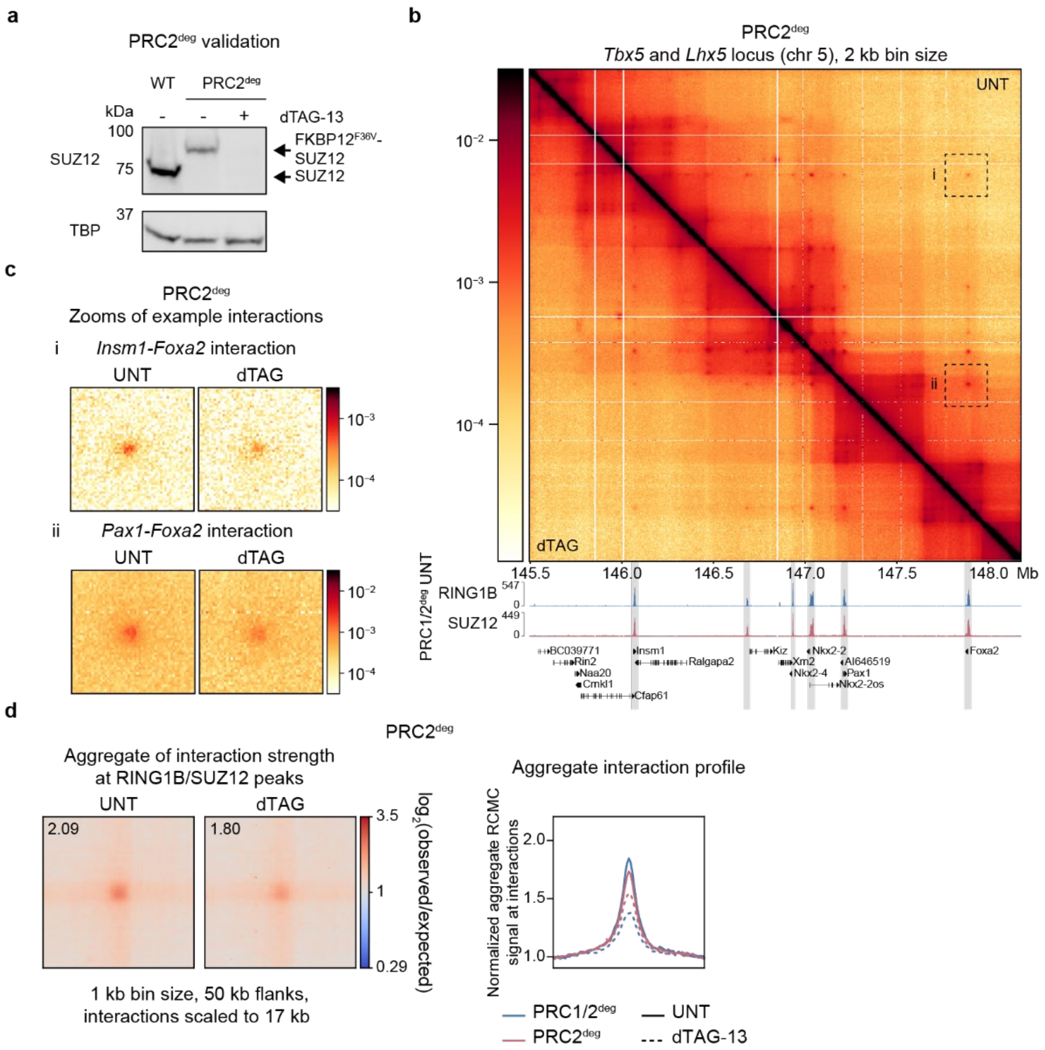


**Supplementary Figure 5. 4 h PRC2 depletion weakens Polycomb-mediated interactions. a,** Western blot validation of SUZ12 tagging and depletion following 4 h dTAG-13 treatment in PRC2$^{deg}$. TBP is included as a loading control. *n* = 1 biological replicate. **b,** RCMC $\log_{10}$-scaled contact maps for PRC2$^{deg}$ showing changes in contact frequency (upper) with corresponding RING1B and SUZ12 ChIP-seq in PRC1/2$^{deg}$ UNT illustrating the locations of Polycomb-occupied regions. Data shown are merges of *n* = 2 (RCMC) or *n* = 3 (ChIP-seq) biological replicates. **c,** Zooms of interactions from RCMC contact map regions labeled i and ii in **b** in untreated and dTAG-13 treatment, labeled according to the genes at the interacting loci. Contact map zooms are plotted at 2 kb bin size. **d,** Left panel: aggregate pileups of interactions ($\log_2$(observed/expected)) between RING1B/SUZ12 co-occupied sites in SUZ12$^{deg}$ with or without 4 h dTAG-13 treatment. Sites are scaled to 17 kb with 50 kb flanks, with data plotted at 1 kb bin size. Values in the upper left corner of each plot indicate the mean value of the central 3x3 pixels of the pileup. Right panel: a 1D average plot across the 21 middle diagonals of the aggregate pileups in the left panel and **Fig. 1f** for PRC1/2$^{deg}$, normalized to the mean of the 10 bins on the left- and right-most sides.

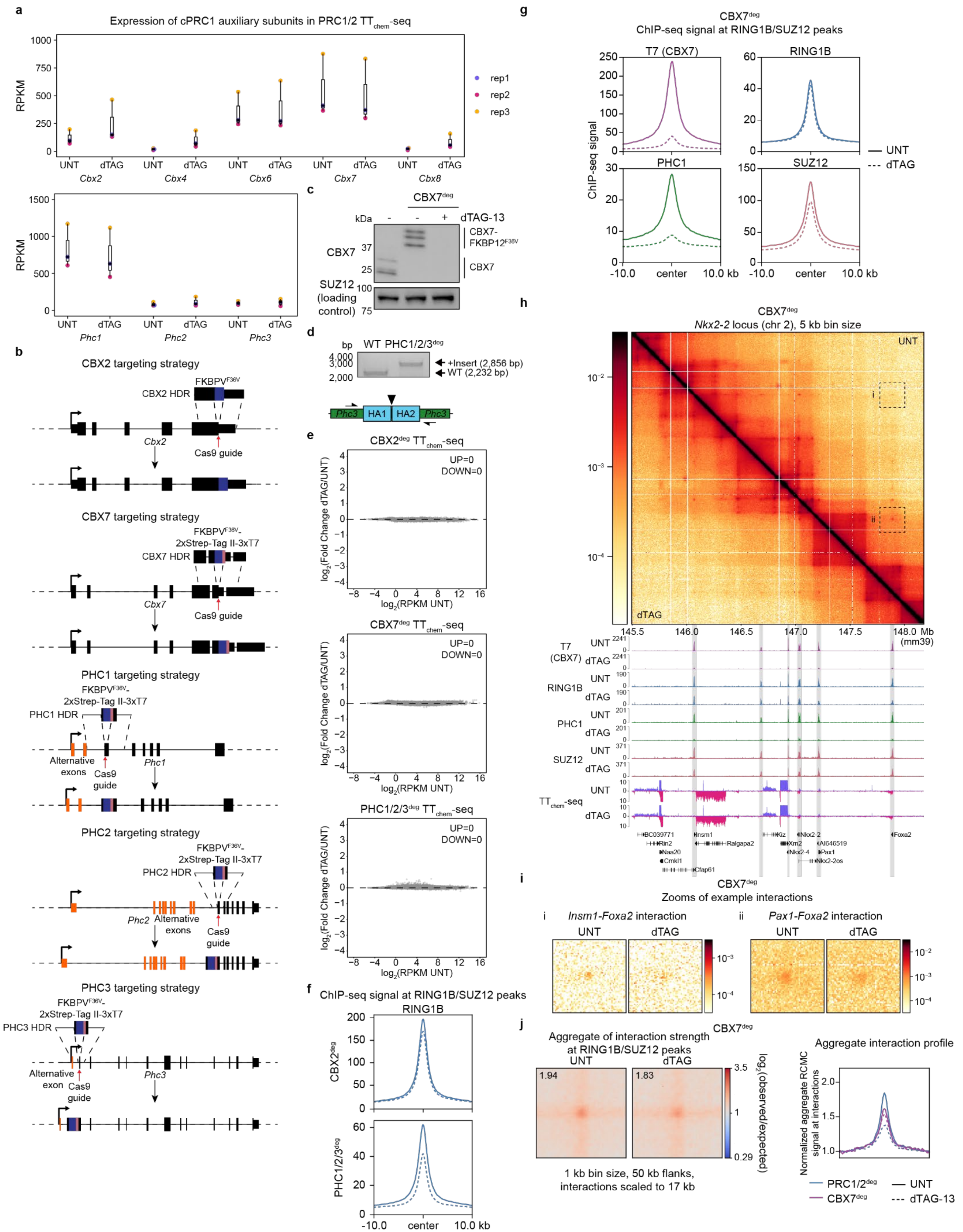


**Supplementary Figure 6. CBX7 depletion displaces cPRC1 from chromatin but does not fully remove Polycomb-mediated interactions. a,** Plot comparing RPKM values for cPRC1 subunits in PRC1/2$^{deg}$ cells with or without 4 h dTAG-13 treatment for *n* = 3 replicates. For boxplots, boxes represent the interquartile range (IQR), the middle line corresponds to the median, and whiskers extend to the largest and smallest values no more than 1.5 x IQR from the box. **b,** A schematic illustrating the genome editing strategy for inserting the $FKBPV^{F36V}$ degron tag into the *Cbx2*, *Cbx7*, and *Phc1/2/3* genes. For *Phc* genes, tags are introduced within the first common exon following multiple alternative exons (orange) to ensure tagging of all isoforms. Narrow boxes indicate untranslated sequences, while wider boxes are translated exons. **c,** Western blot validation of CBX7 tagging and depletion following 4 h dTAG-13 treatment in $CBX7^{deg}$. SUZ12 is included as a loading control. *n* = 1 biological replicate. **d,** DNA agarose gel image showing homozygous insertion of the degron tag in the *Phc3* gene. *n* = 1 replicate. **e,** MA plots of $log_2$(fold change) in nascent gene expression measured by $TT_{chem}$-seq from UNT to dTAG in $CBX2^{deg}$, $CBX7^{deg}$, and PHC1/2/$3^{deg}$. Numbers of significantly differentially up- (UP) or down-changed (DOWN) genes are shown within each plot, and genes which are significantly changed in expression ($log_2$(fold change) > 1.5 and $p_{adj} < 0.05$) are marked with red dots while unchanged genes are shown as grey dots. Data are derived from *n* = 3 biological replicates. **f,** Metaplots showing change of ChIP-seq signal for RING1B at RING1B/SUZ12 co-occupied peaks following 4 h dTAG-13 treatment in $CBX2^{deg}$ and PHC1/2/$3^{deg}$. Data are derived from *n* = 3 biological replicates. **g,** Metaplots showing change of ChIP-seq signal for T7 (CBX7), RING1B, PHC1, and SUZ12 at RING1B/SUZ12 co-occupied peaks following 4 h dTAG-13 treatment in $CBX7^{deg}$. Data are derived from *n* = 3 biological replicates. **h,** RCMC $log_{10}$-scaled contact maps for $CBX7^{deg}$ showing changes in contact frequency (upper) with corresponding ChIP-seq for T7 (CBX7), RING1B, PHC1, and SUZ12, showing decreases in Polycomb subunit occupancy, along with $TT_{chem}$-seq (lower). Data shown are merges of *n* = 2 (RCMC) or *n* = 3 (ChIP-seq and $TT_{chem}$-seq) biological replicates. **i,** Zooms of interactions from RCMC contact map regions labeled i and ii in **h** in untreated and dTAG-13 treatment, labeled according to the genes at the interacting loci. Contact map zooms are plotted at 2 kb bin size. **j,** Left panel: aggregate pileups of interactions (left) ($log_2$(observed/expected)) between RING1B/SUZ12 co-occupied sites in $CBX7^{deg}$ with or without 4 h dTAG-13 treatment. Sites are scaled to 17 kb with 50 kb flanks, with data plotted at 1 kb bin size. Values in the upper left corner of each plot indicate the mean value of the central 3x3 pixels of the pileup. Right panel: a 1D average plot across the 21 middle diagonals of the aggregate pileups in the left panel and **Fig. 1f** for PRC1/$2^{deg}$, normalized to the mean of the 10 bins on the left- and right-most sides.

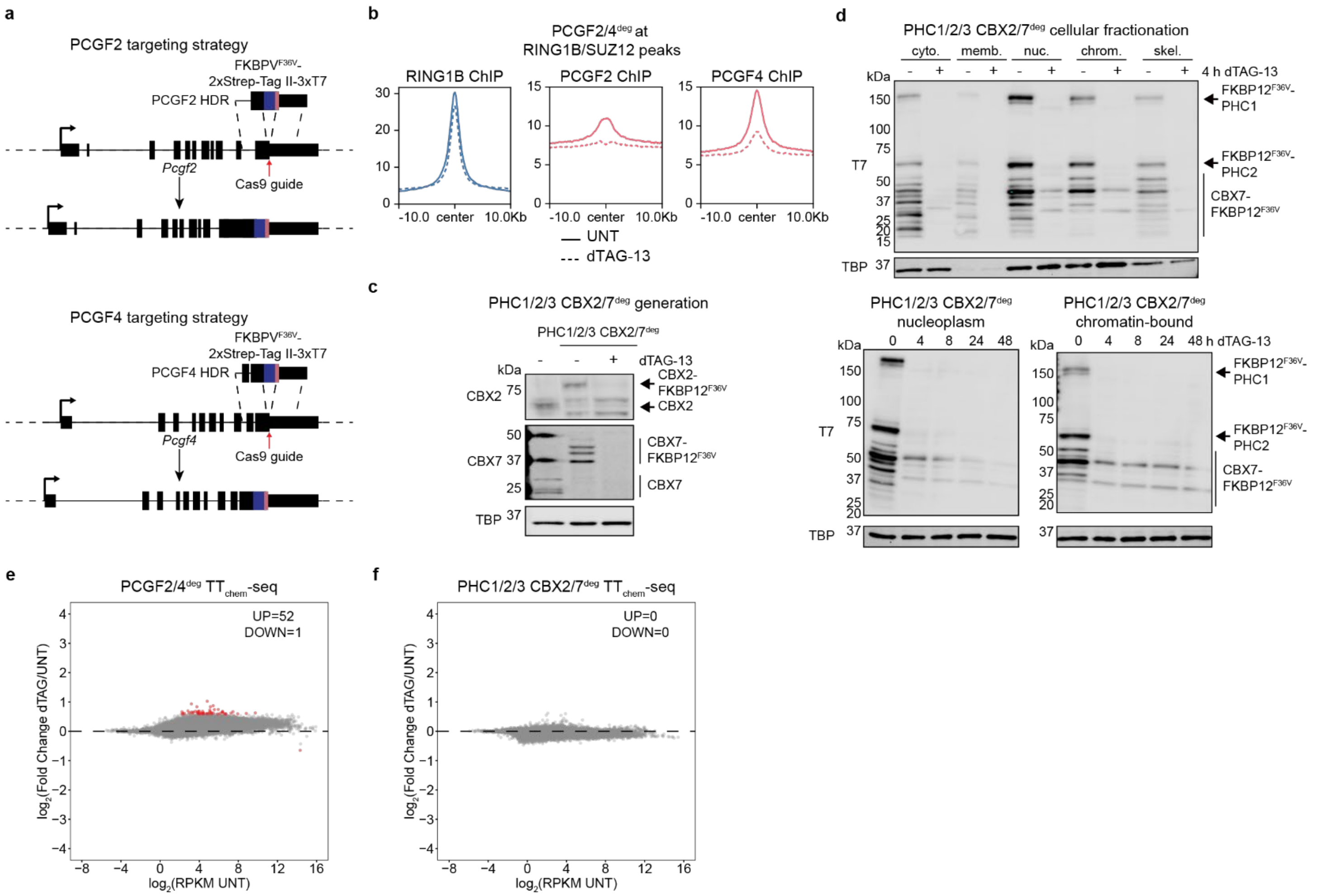


**Supplementary Figure 7. Validation of cPRC1 depletion cell lines. a,** A schematic illustrating the genome editing strategy for inserting the FKBPV$^{F36V}$ degron tag into the *Pcgf2* and *Pcgf4* genes. Narrow boxes indicate untranslated sequences, while wider boxes are translated exons. **b,** Metaplots showing change of ChIP-seq signal for RING1B, PCGF2, and PCGF4 at RING1B/SUZ12 co-occupied peaks following 4 h dTAG-13 treatment in PCGF2/4$^{deg}$. Data are derived from $n$ = 3 biological replicates. **c,** Western blot validation of CBX2 and CBX7 tagging and depletion following 4 h dTAG-13 treatment in the PHC1/2/3$^{deg}$ cell line to generate a PHC1/2/3 CBX2/7$^{deg}$ line. TBP is included as a loading control. $n$ = 1 biological replicate. **d,** Upper panel: western blots of cellular fractionation of PHC1/2/3 CBX2/7$^{deg}$ cells with or without 4 h dTAG-13 treatment. Cyto.: cytosol; memb.: membrane-bound; nuc.: nucleoplasm; chrom.: chromatin-bound; skel.: cytoskeleton. Lower panel: western blot of degron timecourses of dTAG-13 treatment up to 48 h of PHC1/2/3 CBX2/7$^{deg}$ cells for the nucleoplasmic and chromatin-bound fractions. TBP is included as a loading control. $n$ = 1 biological replicate. **e-f,** MA plots of $\log_2$(fold change) in nascent gene expression measured by TT$_{chem}$-seq from UNT to dTAG in PCGF2/4$^{deg}$ (**e**) and PHC1/2/3 CBX2/7$^{deg}$ (**f**). Numbers of significantly differentially up- (UP) or down-changed (DOWN) genes are shown within each plot, and genes which are significantly changed in expression ($\log_2$(fold change) > 1.5 and $p_{adj}$ < 0.05) are marked with red dots while unchanged genes are shown as grey dots. Data are derived from $n$ = 3 biological replicates.

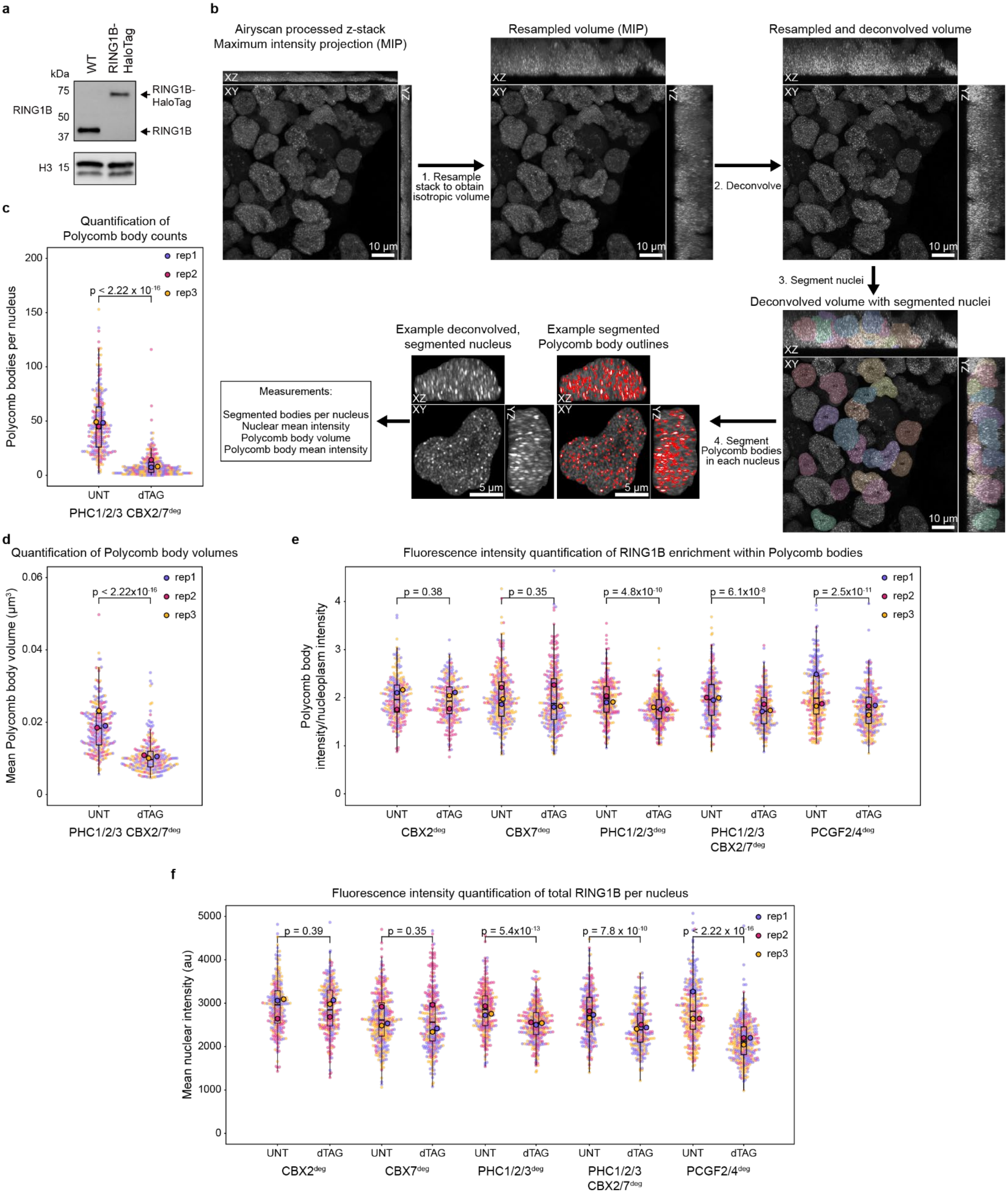


**Supplementary Figure 8. Quantification of Polycomb bodies from live-cell imaging. a,** Western blot validation of RING1B homozygous HaloTag insertion in the parental cell line. Histone H3 is included as a loading control. $n$ = 1 biological replicate. **b,** A schematic of the processing pipeline for analysis of Polycomb bodies based on imaging of RING1B-HaloTag. **c-d,** Plots showing the distribution of the number of Polycomb bodies detected per nucleus (**c**) and mean Polycomb body volume in each nucleus (**d**) for PHC1/2/3 CBX2/7$^{deg}$ nuclei. **e-f,** Plots showing the distributions of Polycomb body mean intensity/nucleoplasm mean intensity in each nucleus (**e**) and mean nuclear intensity of RING1B-HaloTag representing total PRC1 levels in CBX2$^{deg}$, CBX7$^{deg}$, PHC1/2/3$^{deg}$, PHC1/2/3 CBX2/7$^{deg}$, and PCGF2/4$^{deg}$ nuclei. For **c-f**, individual values for each nucleus are colored by treatment replicate (small circles) and are overlaid with a boxplot of the distribution of the individual nuclei and the means of all nuclei in each of three replicates of dTAG-13 treatment (large, outlined circles). For boxplots, boxes represent the interquartile range (IQR), the middle line corresponds to the median, and whiskers extend to the largest and smallest values no more than 1.5 x IQR from the box. Number of nuclei: CBX2$^{deg}$ UNT $n$ = 252; CBX2$^{deg}$ dTAG $n$ = 256; CBX7$^{deg}$ UNT $n$ = 343; CBX7$^{deg}$ dTAG $n$ = 303; PHC1/2/3$^{deg}$ UNT $n$ = 265; PHC1/2/3$^{deg}$ dTAG $n$ = 307; PHC1/2/3 CBX2/7$^{deg}$ UNT $n$ = 257; PHC1/2/3 CBX2/7$^{deg}$ dTAG $n$ = 260; PCGF2/4$^{deg}$ UNT $n$ = 303; PCGF2/4$^{deg}$ dTAG $n$ = 325. Indicated $p$-values were calculated using the Mann-Whitney U Test on per-nucleus values.

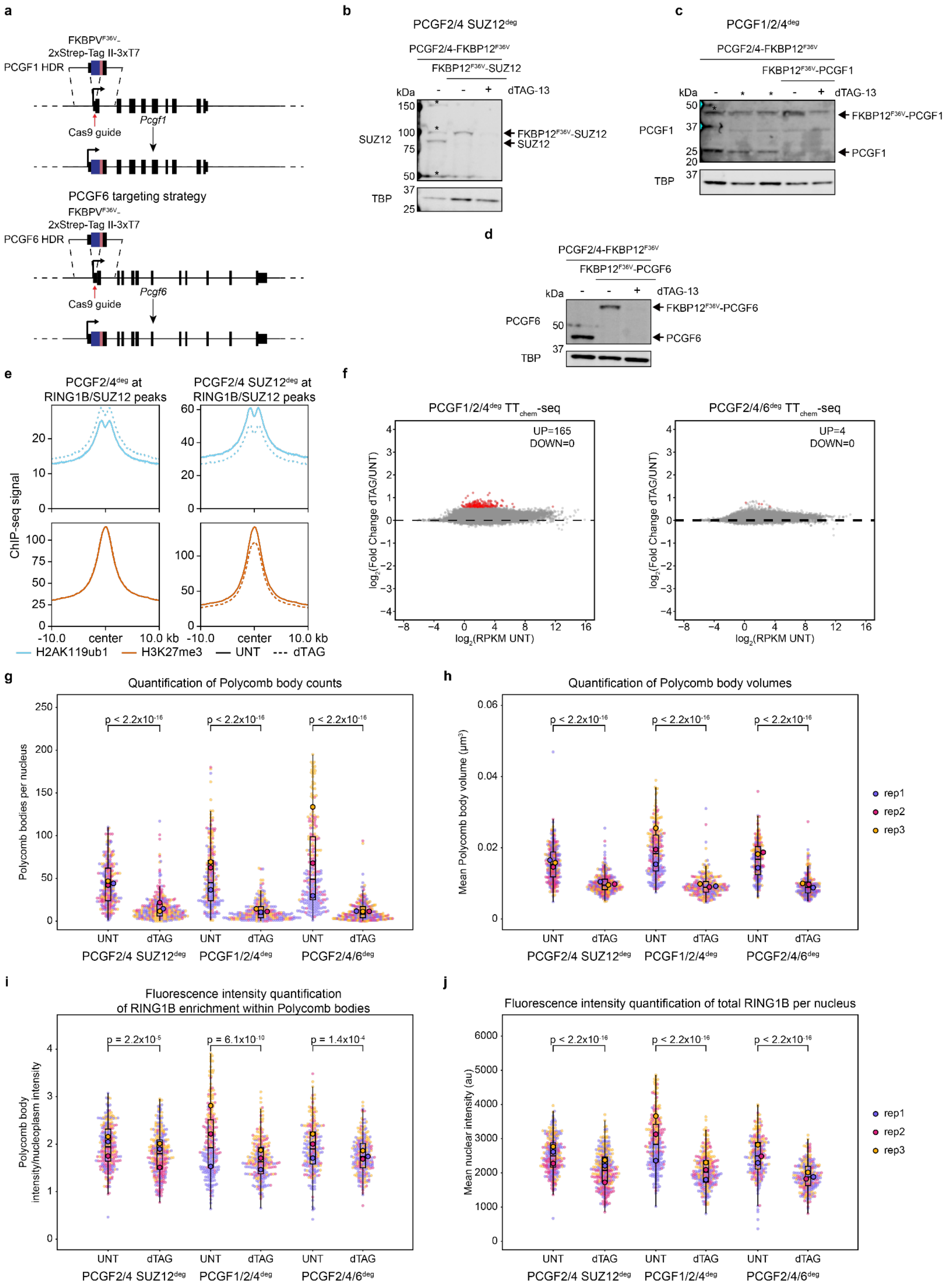
a
FKBPV^F36V-
2xStrep-Tag II-3xT7
PCGF1 HDR
Cas9 guide
Pcgf1
PCGF6 targeting strategy
FKBPV^F36V-
2xStrep-Tag II-3xT7
PCGF6 HDR
Cas9 guide
Pcgf6
b
PCGF2/4 SUZ12^deg
PCGF2/4-FKBP12^F36V
FKBP12^F36V-SUZ12
kDa
dTAG-13
SUZ12
TBP
FKBP12^F36V-SUZ12
SUZ12
c
PCGF1/2/4^deg
PCGF2/4-FKBP12^F36V
FKBP12^F36V-PCGF1
kDa
dTAG-13
PCGF1
TBP
FKBP12^F36V-PCGF1
PCGF1
d
PCGF2/4-FKBP12^F36V
FKBP12^F36V-PCGF6
kDa
dTAG-13
PCGF6
TBP
FKBP12^F36V-PCGF6
PCGF6
e
PCGF2/4^deg at RING1B/SUZ12 peaks
PCGF2/4 SUZ12^deg at RING1B/SUZ12 peaks
ChIP-seq signal
-10.0
center
10.0 kb
H2AK119ub1
H3K27me3
UNT
dTAG
f
PCGF1/2/4^deg TT_chem-seq
UP=165
DOWN=0
PCGF2/4/6^deg TT_chem-seq
UP=4
DOWN=0
log2(Fold Change dTAG/UNT)
log2(RPKM UNT)
g
Quantification of Polycomb body counts
Polycomb bodies per nucleus
p < 2.2x10^-16
UNT
dTAG
PCGF2/4 SUZ12^deg
PCGF1/2/4^deg
PCGF2/4/6^deg
h
Quantification of Polycomb body volumes
Mean Polycomb body volume (µm³)
p < 2.2x10^-16
rep1
rep2
rep3
i
Fluorescence intensity quantification of RING1B enrichment within Polycomb bodies
Polycomb body intensity/nucleoplasm intensity
p = 2.2x10^-5
p = 6.1x10^-10
p = 1.4x10^-4
j
Fluorescence intensity quantification of total RING1B per nucleus
Mean nuclear intensity (au)
p < 2.2x10^-16
rep1
rep2
rep3

**Supplementary Figure 9. Generation and characterization of combined Polycomb degrons. a,** A schematic illustrating the genome editing strategy for inserting the FKBPV$^{F36V}$ degron tag into the *Pcgf1* and *Pcgf6* genes. Narrow boxes indicate untranslated sequences, while wider boxes are translated exons. **b-d,** Western blot validation of SUZ12, PCGF1, and PCGF6 tagging and depletion following 4 h dTAG-13 treatment in the PCGF2/4$^{deg}$ cell line to generate PCGF2/4 SUZ12$^{deg}$, PCGF1/2/4$^{deg}$, and PCGF2/4/6$^{deg}$ cell lines. TBP is included as a loading control. Asterisks on blots indicate spillover of ladder (**b**) or non-specific bands (**c**). Lanes marked with asterisks indicate an unused heterozygous clone. *n* = 1 for all blots. **e,** Metaplots comparing changes in ChIP-seq signal of Polycomb-mediated histone modifications at RING1B/SUZ12 co-occupied peaks following 4 h dTAG-13 treatment in PCGF2/4$^{deg}$ and PCGF2/4 SUZ12$^{deg}$. Data are derived from *n* = 3 biological replicates. **f,** MA plots of $\log_2$(fold change) in nascent gene expression measured by $TT_{chem}$-seq from UNT to dTAG in PCGF1/2/4$^{deg}$ (left) and PCGF2/4/6$^{deg}$ (right). Numbers of significantly differentially up- (UP) or down-changed (DOWN) genes are shown within each plot, and genes which are significantly changed in expression ($\log_2$(fold change) > 1.5 and $p_{adj} < 0.05$) are marked with red dots while unchanged genes are shown as grey dots. Data are derived from *n* = 3 biological replicates. **g-j,** Plots showing the distribution of the number of Polycomb bodies detected per nucleus (**g**), mean Polycomb body volume in each nucleus (**h**), Polycomb body mean intensity/nucleoplasm mean intensity in each nucleus (**i**), and mean nuclear intensity of RING1B-HaloTag representing total PRC1 levels (**j**) for PCGF2/4 SUZ12$^{deg}$, PCGF1/2/4$^{deg}$, and PCGF2/4/6$^{deg}$ nuclei. Individual values for each nucleus are colored by treatment replicate (small circles) and are overlaid with a boxplot of the distribution of the individual nuclei and the means of all nuclei in each of three replicates of dTAG-13 treatment (large, outlined circles). For boxplots, boxes represent the interquartile range (IQR), the middle line corresponds to the median, and whiskers extend to the largest and smallest values no more than 1.5 x IQR from the box. Number of nuclei: PCGF2/4 SUZ12$^{deg}$ UNT *n* = 279; PCGF2/4 SUZ12$^{deg}$ dTAG *n* = 310; PCGF1/2/4$^{deg}$ UNT *n* = 254; PCGF1/2/4$^{deg}$ dTAG *n* = 266; PCGF2/4/6$^{deg}$ UNT *n* = 237; PCGF2/4/6$^{deg}$ dTAG *n* = 253. Indicated *p*-values were calculated using the Mann-Whitney U Test on per-nucleus values.

### METHODS

#### Cell culture

E14-derived mESCs were cultured in Knockout DMEM (ThermoFisher, #10829018) supplemented with 15% fetal bovine serum (FBS; Avantor, #89510-186, lot #266M25), 1,000 U/mL LIF (Cell Guidance Systems, #GFM200), 1x MEM non-essential amino acids (ThermoFisher, #11140050), 1x penicillin/streptomycin (ThermoFisher, #15070063), 1x GlutaMAX (ThermoFisher, #35050061) and 100 µM β-mercaptoethanol (ThermoFisher, #31350010) on plates coated with 0.1% gelatin (Sigma-Aldrich, #G1890-500G) at 37 °C with 5% $CO_2$. For passaging and cell collection, cells were washed once with PBS (ThermoFisher, #10010049), and then incubated with TrypLE Express Enzyme (ThermoFisher, #12605036) for 3 minutes at 37 °C. Cells were resuspended in growth medium before passaging to a new plate or pelleting at 300*g* for 3 minutes. Unless stated otherwise, dTAG-13 treatment was performed by replacing medium with either fresh culture medium (UNT) or medium supplemented with 100 nM dTAG-13 (dTAG) (Fisher Scientific, #66-055). Both treated and untreated cells were replaced in the incubator for 4 hours of treatment prior to performing experiments.

*Drosophila melanogaster* Sg4 cells (RRID:CVCL_Z707; https://dgrc.bio.indiana.edu//stock/RRID:CVCL_Z707)[3] for spike-in were grown at 25 °C in Schneider's *Drosophila* medium (ThermoFisher, #21720024), supplemented with 1X penicillin/streptomycin (ThermoFisher, #15070063), and 10% FBS (Avantor, #89510-186, lot #266M25) which was heat inactivated at 56 °C for 30 minutes prior to cooling to 25 °C for use or freezing at -20 °C. Sg4 cells were detached by pipetting for passaging and cell collection. Cells were plated in 1:4 or 1:5 ratios of cultured to fresh medium. To pellet cells for cell collection, Sg4 cells were centrifuged at 500*g* for 5 minutes.

HEK293T cells for spike-in were grown in high glucose DMEM (ThermoFisher, #31053036) supplemented with 10% FBS (Avantor, #89510-186, lot #266M25), 1x MEM non-essential amino acids (ThermoFisher, #11140050), 1x penicillin/streptomycin (ThermoFisher, #15070063), 1x GlutaMAX (ThermoFisher, #35050061) and 50 µM β-mercaptoethanol (ThermoFisher, #31350010) at 37 °C with 5% $CO_2$. For passaging and cell collection of HEK293T cells, cells were washed once with PBS (ThermoFisher, #10010049), and then incubated with Tryple (ThermoFisher, #12605036) for 3 minutes at 37 °C. Cells were resuspended in growth medium before passaging to a new plate or pelleting at 300*g* for 3 minutes.

#### Genome editing

All cell lines used in this study were derived from an E14 mESC line with TetON3G[4] inserted at the *Rosa26* locus and mCherry-NGN2 under the control of the tight TRE promoter with a bGH poly(A) signal inserted at the TIGRE locus[5,6]. All degron cell lines with the exception of RING1A/B degron lines (PRC1$^{deg}$ and PRC1/2$^{deg}$), were further derived from a cell line in which RING1B was tagged at the C-terminus with HaloTag as previously described[7].

FKBP12$^{F36V}$ degron tags for the dTAG system[8] were added either to the N- or C-terminus of proteins depending on their structure and function, and in some cases including epitope tags to enable ChIP-seq. For RING1A, RING1B, and SUZ12, FKBP12$^{F36V}$ was added at the N-terminus with a 22 amino acid glycine/serine (GS) linker (GGSGGSGSGGGGGSGGGGSGGG). CBX2 was tagged at the C-terminus with GS-linker-FKBP12$^{F36V}$. PCGF2-GS-linker-FKBP12$^{F36V}$-3XT7 cell lines were generated as previously described[9]. PCGF4 and CBX7 were C-terminally tagged with GS-linker-FKBP12$^{F36V}$-3XT7 using the same tag sequence as PCGF2, while PCGF1 and PCGF6 were tagged at the N-terminus using 3XT7-FKBP12$^{F36V}$-GS linker and 3XFLAG-FKBP12$^{F36V}$-GS-linker sequences, respectively. Since C-terminal tagging has previously been shown to disrupt the function of the SAM domain of PHC proteins, and because all three *Phc1*, *Phc2*, and *Phc3* genes have complex alternative splicing at the 5' end of their coding regions, a GS-linker-FKBP12$^{F36V}$-3XT7-GS-linker sequence was inserted at the beginning of the 5'-most common exon for all isoforms (Fig S6b) after confirming using InterPro[10] that this sequence was otherwise unstructured.

For insertion of arrays for locus imaging, the 982-bp ANCHOR3 sequence from NeoVirTech[11] and the 1331-bp synBsr1 sequence[12] were inserted in the PRC1/2$^{deg}$, PCGF2/4$^{deg}$, and PHC1/2/3$^{deg}$ degron lines generated above. ANCHOR3 was inserted 833 bp upstream of the transcription start site (TSS) of *Tbx5*, and synBsr1 was either inserted 1843 bp upstream of *Lhx5* TSS or at position chr5: 119,369,198, which is an empty region 601,043 bp upstream of the *Tbx5* gene. After confirming homozygous insertion of the arrays, an OR3[11] protein C-terminally tagged with two copies of mStayGold(J)[13,14] and SV40 nuclear localization signal (OR3-2xmStayGold(J)-3xNLS) and a synBsr1-ParB[12] protein C-terminally tagged with two copies of the mScarlet3[15] and SV40 nuclear localization signal (synBsr1-2xmScarlet3-NLS) were integrated using the PiggyBac transposase system. Cells were transfected with the above fluorescently fused binding proteins and a plasmid encoding Super PiggyBac Transposase at a ratio of 4:1 using Lipofectamine 3000. Fluorescent cells were initially selected using fluorescence activated cell sorting two weeks after transfection. Following the initial sort, cells were sorted twice more, once after one week after the first sort and another the following week. For the last sort, cells were divided into three pools based on their fluorescence levels and each pool was visually inspected using a Zeiss LSM900 Airyscan microscope to determine the pool with the most cells with visible OR3 and synBsr1 foci. Then, 6,000 cells for each cell line from the pool with the most optimal fluorescence were plated on 15-cm dishes and allowed to grow to individual colonies. 96 clones from each cell line were picked, plated on a Revvity glass-bottomed 96-well plate, and screened using an Opera Phenix High-Content screening system. Two to three clones with clearly visible OR3 and synBsr1 dots for each cell line were pooled for imaging experiments.

All mESC lines expressing endogenously tagged proteins in this study were generated using CRISPR-Cas9-mediated genome editing. Homology-directed repair (HDR) templates were cloned containing homology sequences matching the genomic (mm39) sequence ~1 kb up- and downstream of the insertion site (primer sequences for cloning homology arms are listed in Table S3), flanking the desired insertion. SpCas9-compatible guides were selected using CRISPOR[16] to identify appropriate Gn19 guide sequences (Table S4), and were cloned into the pSpCas9(BB)-2A-Puro plasmid (PX459)[17]. Guide plasmids and HDR templates were transfected into cells at 70-80% confluency in a 6-well plate using Lipofectamine 3000 (ThermoFisher, #L3000015) according to the manufacturer's protocol, leaving the Lipofectamine mix on the cells overnight. The following morning, cells were passaged 1:20 to 10 cm plates. After attaching, cells were selected for transfection of the guide plasmid using 1 µg/mL puromycin (Sigma-Aldrich, #P8833) for 48 hours. Following puromycin removal, cells were allowed to grow into clonal colonies before being picked. Individual clones were screened for homozygous editing by extraction of gDNA followed by PCR (screening primers can be found in Table S3), followed by western blotting for protein tagging.

#### Extracts

gDNA was prepared from cells for screening by freezing attached cells on the plate at -80 °C for at least 30 minutes. Cells were then thawed at room temperature and resuspended in squishing buffer (10 mM Tris HCl pH 8, 1 mM EDTA, 25 mM NaCl, 200 µg/mL proteinase K (Viagen Biotech, #501-PK)) with scraping with a pipette tip. Resuspended cells were lysed by incubation at 65 °C for 1 hour. The resulting lysate was used for screening PCRs.

Nuclear extracts were prepared from mESCs which had been collected by pelleting and frozen at -80 °C. Pellets were thawed on ice, and cells resuspended in 10 volumes buffer A (10 mM HEPES pH 7.9, 1.5 mM $MgCl_2$, 10 mM KCl, 0.5 mM PMSF, and 1X cOmplete protease inhibitors EDTA-free (Sigma-Aldrich, #11873580001)) and incubated on ice for 10 minutes. Cells were pelleted at 1500*g* for 5 minutes, 4 °C, and then resuspended in three volumes of buffer A supplemented with 0.1% IGEPAL CA-630 (Sigma-Aldrich, #I8896-50ML), followed by inversion 10 times. Nuclei were pelleted as before, and then resuspended in buffer C (5 mM HEPES pH 7.9, 26% glycerol, 1.5 mM $MgCl_2$, 0.2 mM EDTA, 1X cOmplete protease inhibitors, 0.5 mM DTT, and 250 mM NaCl). The volume of the resuspended nuclei was measured by pipetting, and NaCl added dropwise to bring the final concentration to 400 mM. Nuclei were then incubated for 1 hour on ice with occasional agitation, and then pelleted at 16,000*g*. The resulting supernatant was taken as the nuclear extract.

Histone extracts were prepared from mESCs which had been collected by pelleting and frozen at -80 °C. For quantification, pellets were collected from three separate treatments on different days. After thawing on ice, cells were resuspended in RSB (10 mM Tris HCl pH 7.4, 10 mM NaCl, 3 mM $MgCl_2$, 20 mM N-ethylmaleimide (NEM, Fisher Scientific, #AA4052614)). Cells were then pelleted at 300*g*, 4 °C for 5 minutes, and then resuspended in RSB supplemented with 0.5% IGEPAL CA-630 (Sigma-Aldrich, #I8896-50ML) and incubated on ice for 10 minutes. Nuclei were pelleted at 500*g*, 4 °C for 5 minutes, and then resuspended in 5 mM $MgCl_2$ with 20 mM NEM. 1 volume of 0.8 M HCl was added to the nuclei suspension, and the nuclei were incubated on ice for 20 minutes. Nuclei were pelleted at 16,000*g*, 4 °C for 20 minutes, and the resulting supernatant transferred to a new tube. Trichloroacetic acid (Sigma-Aldrich, #T0699-100ML) was added to 25% v/v and the mixture incubated on ice for 30 minutes to precipitate proteins. Histones were pelleted by centrifugation at 16,000*g*, 4 °C for 15 minutes, and pellets were washed with ice cold acetone. Histones were pelleted again as before and the acetone wash repeated. After removal of acetone from the second wash, pellets were air-dried to remove remaining acetone, before resuspending the histone pellets in 1X SDS loading buffer (2% SDS, 0.1 M Tris pH 6.8, 0.1 M DTT, 10% glycerol, bromophenol blue) and heating at 90 °C for 5 minutes to redissolve histones.

#### Western blotting

For western blotting, nuclear extracts were run on 4-12% gradient Tris-glycine SDS-PAGE gels (ThermoFisher, #XP04122BOX) or histone extracts were run on 15% Tris-glycine SDS-PAGE gels (ThermoFisher, #XP00145BOX) with Precision Plus Protein dual color standards (Bio-Rad, #1610374S). Gels were run in 1X Tris-glycine-SDS buffer (Bio-Rad, #1610732) at 200 V (nuclear extracts) or 150 V (histone extracts). Proteins were transferred to 0.2 µm nitrocellulose membrane (Bio-Rad, #1620112) using an eBlot L1 wet transfer system (GenScript, #L00686) on the standard (nuclear extract, 16.5 min) or short (histone extract, 11.16 min) setting. Membranes were blocked with 5% w/v milk (Santa Cruz Biotechnology, #sc-2325) in PBS with 0.1% Tween 20 (PBS-T) for 30 minutes at room temperature. Blots were probed with primary antibodies (primary antibodies and dilutions are listed in Table S5) diluted in 5% milk in PBS-T by overnight incubation at 4 °C. Blots were rinsed once with PBS, and then washed three times for 5 minutes each with PBS-T. Blots were then probed with either HRP- or fluorescently-conjugated secondary antibodies (secondary antibodies and dilutions are listed in Table S5) for 1 hour at room temperature. Following secondary antibody incubation, blots were rinsed once with PBS, washed three times for 5 minutes each with PBS-T, followed by a final PBS rinse. Blots probed with HRP-conjugated secondary antibodies were imaged using Clarity Max Western ECL Substrate (Bio-Rad, #1705062), and both ECL and fluorescent images were taken using a ChemiDoc (Bio-Rad, #12003154). Gel band quantification was performed in ImageStudio (v5.5, LICORbio) software. Blots for quantification were performed three times with all quantified samples to account for technical variation.

#### Crosslinked ChIP-seq

Crosslinked calibrated ChIP-seq[18–20] was performed as described previously[21]. mESC samples were prepared from three independent treatments on different days. mESCs with or without dTAG-13 treatment, or HEK293T cells for spike-in, were counted, and $5x10^7$ cells were pelleted and resuspended in PBS. Cells were crosslinked first by addition of disuccinimidyl glutarate (ProteoChem, #c1104-100mg) to 2 mM and incubation at RT for 45 minutes, followed by addition of formaldehyde (ThermoFisher, #28908) to 1% and incubation at RT for 10 minutes. Crosslinking was quenched by addition of glycine to 125 mM. Crosslinked cells were washed once with ice-cold PBS, and then pelleted and snap frozen in liquid nitrogen. Prior to snap freezing, HEK293T cells were aliquoted into batches of $2x10^6$ cells. Crosslinked mESCs were thawed on ice and then resuspended in LB1 (10% glycerol, 50 mM HEPES-KOH pH 7.9, 140 mM NaCl, 1 mM EDTA, 0.5% IGEPAL CA-630 (Sigma-Aldrich, #I8896-50ML), 0.25% Triton X-100, 1X cOmplete protease inhibitors EDTA-free (Roche, #04693159001) and incubated on ice for 10 minutes. Nuclei were pelleted by centrifugation at 1000*g*, 4 °C for 5 minutes. Nuclei were washed with LB2 (10 mM Tris-HCl pH 8.0, 200 mM NaCl, 1 mM EDTA, 0.5 mM EGTA, 1X cOmplete protease inhibitors EDTA-free) for 5 minutes on ice before pelleting as before. Nuclei were then resuspended in LB3 (10 mM Tris-HCl pH 8.0, 100 mM NaCl, 1 mM EDTA, 0.5 mM EGTA, 0.1% sodium deoxycholate, 0.5% N-lauroylsarcosine, 1X cOmplete protease inhibitors EDTA-free) and transferred to milliTUBE 1 mL with AFA fiber (Covaris, #520135) sonication tubes. Chromatin was sonicated on a Covaris E220evolution focused-ultrasonicator with the following settings: 17.5 minute sonication time, 140 W peak incident power, 5% duty factor, 200 cycles/burst, 4 °C. Following sonication, Triton X-100 was added to a concentration of 1%, cell debris was pelleted, and the supernatant was taken as the chromatin. Chromatin was quantified following 1:10 dilution in 0.1 M NaOH using a Nanodrop instrument.

20 µL of Protein A Dynabeads (ThermoFisher, #10001D) per pulldown were washed once with ChIP dilution buffer (1 mM EDTA, 20 mM Tris-HCl pH 8.0, 150 mM NaCl, 1% Triton X-100) and then incubated with blocking buffer (ChIP dilution buffer with 0.2 mg/mL BSA (Sigma-Aldrich, #B8667-1.25ML), 50 µg/mL yeast tRNA (ThermoFisher, #AM7119)) at 4 °C for 1 hour. After blocking, beads were washed twice more with ChIP dilution buffer. 150 µg of chromatin was diluted 1:10 in ChIP dilution buffer for each pulldown. Diluted chromatin was precleared by addition of 10 µL of blocked beads per pulldown and incubation for 1 hour at 4 °C. Following preclearing, antibody was added according to Table S5, and samples were incubated overnight at 4 °C. 15 µg of precleared chromatin was reserved as input. 10 µL of beads was added to each pulldown to bind antibody, and beads were then washed at 4 °C for 4 minutes each with low salt buffer (0.1% SDS, 1% Triton X-100, 2 mM EDTA, 20 mM Tris-HCl pH 8, 150 mM NaCl), high salt buffer (0.1% SDS, 1% Triton X-100, 2 mM EDTA, 20 mM Tris-HCl pH 8, 500 mM NaCl), LiCl wash buffer (250 mM LiCl, 0.1% IGEPAL CA-630, 25 mM sodium deoxycholate, 5 mM EDTA, 10 mM Tris-HCl pH 8.0), and two washes with 1X Tris-EDTA buffer (Sigma-Aldrich, #93283-100ML). Chromatin was eluted by resuspending beads in elution buffer (1% SDS, 0.1 M $NaHCO_3$) and vortexing for 30 minutes at RT.

DNA was purified from isolated chromatin and from inputs by addition of NaCl to 0.2 M and RNase A to 0.2 mg/mL, and incubating at 65 °C overnight with 1000 rpm shaking. The next morning, proteinase K was added to 0.2 mg/mL, and samples were incubated at 45 °C for 1 hour with shaking. DNA was purified using the Zymo ChIP DNA Clean & Concentrator kit (Zymo Research, #D5205). Purified DNA was sonicated using a Covaris E220evolution focused-ultrasonicator to reduce fragment sizes in microTUBE AFA Fiber tubes (Covaris, #520045) with the following settings: 50 s sonication time, 105 W peak incident power, 5% duty factor, 200 cycles/burst, 4 °C. Libraries for sequencing were then prepared using the NEBNext UltraExpress DNA Library Prep Kit (NEB, #E3325L) according to the manufacturer's protocol and indexed using NEB Multiplex Oligos for Illumina (Dual Index) (NEB, #E7600 and #E7780). Libraries were sequenced on the Illumina NovaSeq X sequencing platform with 50-bp each end paired reads.

#### Native ChIP-seq

Calibrated ChIP-seq[18–20] of histone modifications was performed as previously described[21]. Sg4 *Drosophila melanogaster* cells were prepared in advance for spike-in. Sg4 cells were washed with PBS and aliquoted for later use prior to snap freezing. mESC samples were prepared from three independent treatments on separate days for PCGF2/4$^{deg}$ and PCGF2/4

SUZ12$^{deg}$ cells, and from a single treatment for PRC1/2$^{deg}$ cells. To isolate chromatin, mESCs with or without dTAG-13 treatment were counted, and 5x10$^7$ cells were pelleted and resuspended in PBS. 2x10$^7$ thawed Sg4 cells were added to each sample of mESCs, and the mixture was pelleted. Cells were then resuspended in 1 mL ice cold RSB (10 mM Tris-HCl pH 8.0, 10 mM NaCl, 3 mM $MgCl_2$), before adding 28 mL RSB supplemented with 0.1% IGEPAL CA-630 (Sigma-Aldrich, #I8896-50ML) and 5 mM N-ethylmaleimide (NEM, Fisher Scientific, #AA4052614) and mixing by gentle inversion. Nuclei were pelleted at 1500*g* for 5 minutes at 4 °C, and then resuspended in RSB with 0.25 M sucrose, 3 mM $CaCl_2$, 1X cOmplete protease inhibitors EDTA-free, and 5 mM NEM. Nuclei were pelleted as before and resuspended in 1 mL of the same buffer. Chromatin was digested by addition of 150 U of MNase (Worthington Biochemical. #LS004798) and incubation at 37 °C for 5 minutes with 1,000 rpm shaking. Digestion was stopped by addition of 4 mM EDTA. Nuclei were pelleted at 2,400*g* for 5 minutes at 4 °C, and the supernatant was taken as the first chromatin fraction. Pellets were then resuspended in 300 µL nucleosome release buffer (10 mM Tris-HCl pH 7.5, 10 mM NaCl, 0.2 mM EDTA, 1X cOmplete protease inhibitors EDTA-free, 10 mM NEM) and incubated at 4 °C for 1 hour with mixing. The suspensions were sheared by passing them through a 27G needle 5 times. Cell debris was pelleted at 2,400*g* for 5 minutes at 4 °C, and the supernatant was taken as the second chromatin fraction and combined with the first fraction. Chromatin was then stored at -80 °C until used for pulldowns.

For pulldowns, 100 µL of thawed combined chromatin fractions was diluted 1:10 in native ChIP incubation buffer (70 mM NaCl, 10 mM Tris-HCl pH 7.5, 2 mM $MgCl_2$, 2 mM EDTA, 0.1% Triton X-100, 10 mM NEM, 1X cOmplete protease inhibitors EDTA-free). Any precipitate was removed by centrifugation at 2400*g* for 5 minutes, 4 °C. 100 µL of diluted chromatin was taken as 10% input, and 1 mL of diluted chromatin was used per pulldown and incubated with antibody overnight at 4 °C according to Table S5. For each IP, 20 µL of Protein A Dynabeads (ThermoFisher, #10001D) were incubated in blocking buffer (native ChIP incubation buffer with 1 mg/mL BSA (Sigma-Aldrich, #B8667-1.25ML), 1 mg/mL yeast tRNA (ThermoFisher, #AM7119) for 30 minutes at 4 °C with rotation. Beads were then washed twice with native ChIP incubation buffer, and resuspended in a final volume of 20 µL per pulldown of native ChIP incubation buffer prior to addition to the pulldown. Chromatin/antibody mixes were transferred to a clean tube containing the blocked Dynabeads and incubated with rotation for 1 hour at 4 °C. Beads were then washed four times total with ice cold native ChIP wash buffer (20 mM Tris-HCl pH 7.5, 2 mM EDTA, 125 mM NaCl, 0.1% Triton X-100), followed by one wash with ice cold 1x TE buffer. DNA was eluted from the beads by resuspending in elution buffer (1% SDS, 0.1 M $NaHCO_3$) and vortexing for 30 minutes at RT, followed by purification using the Zymo ChIP DNA Clean & Concentrator kit (Zymo Research, #D5205).

Libraries for sequencing were prepared using the NEBNext UltraExpress DNA Library Prep Kit (NEB, #E3325L) according to the manufacturer's protocol and indexed using NEB Multiplex Oligos for Illumina (Dual Index) (NEB, #E7600 and #E7780). Libraries were sequenced on the Illumina NovaSeq X sequencing platform with 50-bp each end paired reads.

#### TT$_{chem}$-seq

TT$_{chem}$-seq was performed as described previously[22]. For mESCs, treatments were performed on three days, giving three replicates of treatment. Cells were passaged in the morning of the experiment, seeding 5x10$^7$ cells per condition. Once cells attached (2-3 hours), the same volume of fresh culture medium without additions (UNT) or containing 200 nM dTAG-13 (dTAG) was added to the plate, with the treatment carried out for 4 hours. At the end of the treatment time, culture medium was removed and replaced with freshly prepared medium containing 500 µM 4-thiouridine (4SU, Alfa Aesar, #J60679-MD). Cells were incubated with 4SU for 5 minutes to label nascent RNA, and then immediately harvested by removal of medium and direct addition of TRIzol reagent (ThermoFisher, #15596026) to the culture plate. Cells in TRIzol were snap-frozen in liquid nitrogen and stored at -80 °C prior to further processing.

*Drosophila melanogaster* Sg4 cells were prepared as with mESCs for spike-in, but were instead seeded at 1x10$^8$ cells per plate and labeled for 10 minutes. Prior to snap-freezing, Sg4 cell suspensions in TRIzol were aliquoted in 3x10$^7$ cell batches.

TRIzol-suspended samples were thawed at RT, and for each 5x10$^7$ mESC sample, volume equivalent to 1.5x10$^7$ cells of Sg4 suspension in TRIzol was added. The TRIzol mix was transferred to a MaXtract high-density phase-lock tube (Qiagen, #129065). One fifth of the volume of chloroform (Fisher Scientific, #RSOC0020500) was added to the TRIzol suspension and mixed by shaking. After incubation for 3 minutes at RT, organic and aqueous phases were separated by centrifugation for 15 minutes at 2,000*g*, 4 °C. The aqueous phase was then transferred to a fresh tube and RNA precipitated by addition of 1.1 volumes of isopropanol and incubation for 15 minutes at RT. Precipitated RNA was pelleted by centrifugation at 12,000*g* for 20 minutes at 4 °C. Pellets were washed once with 85% ice cold EtOH. All remaining liquid was removed, and the pellet was resuspended in nuclease-free water (ThermoFisher, #AM9937).

Contaminating genomic DNA was removed using the Turbo DNA-free kit (ThermoFisher, #AM1907). Resuspended RNA was incubated in 1X Turbo DNase Buffer with 3 U of Turbo DNase for 30 minutes at 37 °C. The DNase reaction was halted by addition of 0.1 volumes of DNase Inactivation Reagent and incubation for 5 minutes at RT. Inactivation Reagent was pelleted by centrifugation at 16,000*g* for 1.5 minutes, 4 °C, and the supernatant was transferred to a fresh tube. Centrifugation and transfer were repeated to remove any residual Inactivation Reagent. Purified, DNA-free RNA was quantified using the Qubit RNA BR assay kit (ThermoFisher, #Q10210), and 300 µg of RNA was used for the remainder of the protocol.

RNA was diluted to a volume of 100 µL by addition of nuclease-free water, and 20 µL of 1 M NaOH added to fragment RNA by incubation on ice for 20 minutes. Tris-HCl pH 6.8 was added to 400 mM to stop fragmentation, and the solution was applied to a Micro Bio-Spin Column with Bio-Gel P-30 in Tris Buffer (Bio-Rad, #7326224). The solution was passed through the column by centrifugation at 1,000*g* for 4 minutes, RT. The flow-through was then passed through a fresh column for a second cleanup step and diluted to a total volume of 600 µL in nuclease-free water.

4SU was biotinylated by addition of 150 µL of 0.1 mg/mL MTSEA Biotin-XX (Biotium, #90066) in dimethyl formamide and 9 µL of biotin buffer (833 mM Tris-HCl pH 7.4, 83.3 mM EDTA) and incubation for 30 minutes at RT in the dark. Following the biotinylation reaction, the solution was transferred to a phase-lock gel tube (Quantabio, #2302820) and mixed with 1 volume of 25:24:1 phenol:chloroform:isoamyl alcohol (Sigma-Aldrich, #P2069-100ML). Following incubation for 3 minutes at RT, the aqueous and organic phases were separated by centrifugation at 12,000*g* for 5 minutes at 4 °C. The upper, aqueous phase was transferred to a new phase-lock gel tube and mixed again with an equal volume of 25:24:1 phenol:chloroform:isoamyl and centrifuged. After again transferring the aqueous layer to a new tube, RNA was precipitated by addition of 1/10 volume of 5 M NaCl and 1.1 volumes of isopropanol, with incubation for 15 minutes at RT. RNA was pelleted by centrifugation at 20,000*g* for 20 minutes at 4 °C. The RNA pellet was washed with 85% ethanol without resuspending, and the ethanol removed after centrifugation at 20,000*g* for 5 minutes at 4 °C. The RNA pellet was then redissolved in 100 µL of nuclease-free water.

Biotinylated RNA was denatured at 65 °C for 10 minutes followed by cooling on ice for 5 minutes. 200 µL of µMACS streptavidin MicroBeads (from the µMACS Streptavidin Kit, Miltenyi Biotec, #130-133-282) were added to the RNA and incubated with rotation for 15 minutes at RT. A µColumn (from the µMACS Streptavidin Kit, Miltenyi Biotec, #130-133-282) was placed on a µMACS magnetic separator (Miltenyi Biotec, #130-042-602) on a MACS multistand (Miltenyi Biotec, #130-042-303) and prepared by washing with 100 µL nucleic acid equilibration buffer (from the µMACS Streptavidin Kit, Miltenyi Biotec, #130-133-282). The RNA/bead mixture was then applied to the column, and the column was washed three times with 500 µL of pull-down wash buffer (100 mM Tris-HCl pH 7.4, 10 mM EDTA, 1 M NaCl, 0.1% Tween 20) pre-warmed to 55 °C. The column was washed a further three times with 500 µL of RT pull-down wash buffer. RNA was eluted from the streptavidin beads by addition twice of 100 µL freshly prepared elution buffer (100 mM dithiothreitol (DTT)), with each addition separated by 5 minutes.

RNA eluate was cleaned up and concentrated using the RNeasy MinElute Cleanup kit (Qiagen, #74204). To allow more efficient capture of <200 nt RNA fragments, 700 µL of RLT buffer was added to 200 µL of RNA, and 1.05 mL of 100% ethanol added to the mix before applying to the MinElute column. The remainder of the cleanup was performed according to the MinElute Cleanup kit protocol. Libraries for sequencing were prepared using the NEBNext Ultra II Directional RNA Library Prep Kit (NEB, #E7760L) according to the manufacturer's protocol for rRNA depleted FFPE RNA (section 5) and indexed using NEB Multiplex Oligos for Illumina (Dual Index) (NEB, #E7600 and #E7780). Libraries were sequenced on the Illumina NovaSeq X sequencing platform with 50-bp each end paired reads.

#### Region Capture Micro-C

RCMC was performed as described previously[23,24]. Treatments were performed separately on two different days, and $3x10^7$ mESCs were collected for each replicate of each condition. Cells were pelleted at 300*g* for 3 minutes, and then resuspended in 1 mL of PBS per $1x10^6$ cells. Cells were crosslinked with gentle rotation at RT for 35 minutes with 3 mM DSG. After 35 minutes had elapsed, cells were crosslinked for a further 10 minutes with 1% formaldehyde (ThermoFisher, #28908). Crosslinking was quenched with addition of Tris-HCl pH 7.5 to a final concentration of 0.375 M and a further 5 minutes of rotation at RT. Crosslinked cells were pelleted at 1000*g* for 5 minutes at 4 °C and then washed twice with PBS. Following washes, cells were split into $5x10^6$ cell aliquots, pelleted again, and then snap frozen in liquid nitrogen.

The optimal MNase quantity for digesting the crosslinked chromatin was determined for each batch of fixed cells by performing an MNase titration on a single aliquot of $5x10^6$ cells. The aliquot was thawed on ice and then resuspended in buffer MB1 (10 mM Tris-HCl pH 7.5, 50 mM NaCl, 5 mM MgCl2, 1 mM CaCl2, 100 µg/mL BSA (Sigma-Aldrich, #B8667), 1x Protease Inhibitor Cocktail (cOmplete, EDTA-free Protease Inhibitor Cocktail, Sigma-Aldrich, #11873580001), 0.2% NP-40 Alternative (Millipore Sigma #492018)) and incubated on ice for 20 minutes. Nuclei were washed once more with MB1 and then split to $1x10^6$ cell aliquots for MNase digestion. Aliquots were digested using a range of MNase (Worthington Biochemical, #LS004798) quantities between 2 and 20 U by incubating at 37 °C for 20 minutes with 1,000 rpm mixing. MNase digestion was halted by addition of EGTA to 4 mM and heating at 65 °C for 10 minutes. Nuclei were pelleted and resuspended in reverse crosslinking mix (0.8X TE, 1 mg/mL Proteinase K (Viagen Biotech, #501-PK), 1% SDS, 0.1 mg/mL RNase A (ThermoFisher, #EN0531), 200 mM NaCl). Nuclei in reverse crosslinking mix were incubated at 65 °C overnight with 1,000 rpm mixing, and DNA was purified the next morning with the Zymo DNA Clean and Concentrator (DCC-25) kit (Zymo Research, #D4034). Purified DNA was run on a 1% agarose gel, and the optimal MNase determined according to the ratios of mono-, di-, and trinucleosomal band signal.

Micro-C libraries were prepared from $1.5 \times 10^7$ cells for each replicate of each condition, split amongst three $5 \times 10^6$ aliquots. As in the MNase titration, cells were incubated in MB1 for 20 minutes on ice, followed by a single wash and resuspension with MB1. Chromatin was digested using the calculated optimal MNase quantity based on the titration with incubation at 37 °C for 20 minutes. Digestion was quenched by addition of EGTA to 4 mM and heating at 65 °C for 10 minutes. Nuclei were pelleted and washed twice with MB2 (10 mM Tris-HCl pH 7.5, 50 mM NaCl, 10 mM $MgCl_2$, 100 µg/mL BSA) prior to resuspension in 3' end exonuclease reaction mix (1x NEBuffer 2.1 (New England Biolabs, #B7202S), 2 mM ATP (ThermoFisher, #R1441), 5 mM DTT (Sigma-Aldrich, #10197777001), 0.5 U/µL T4 PNK (New England Biolabs, #M0201)) and incubation at 37 °C with constant, 1,000 rpm mixing for 15 minutes. Klenow fragment was then added to 0.5 U/µL, and the samples were incubated again for 15 minutes. Next, biotin labeling reagents (200 µM biotin-14-dATP (Jena Bioscience, #NU-835-BIO14), 200 µM biotin-11-dCTP (Jena Bioscience, NU-809-BIOX), 200 µM dTTP (Jena Bioscience, NU-1004), 200 µM dGTP (Jena Bioscience, #NU-1003), 1x T4 DNA ligase buffer (New England Biolabs, #B0202S), 100 µg/mL BSA) were then added to the samples, and biotin labeling was performed by incubation at 25 °C for 45 minutes with one minute on, three minutes off interval mixing at 1,000 rpm. Labeling was halted by addition of EDTA to 30 mM and incubation with interval mixing at 65 °C for 20 minutes. Nuclei were washed with MB3 (50 mM Tris-HCl pH 7.5, 10 mM $MgCl_2$, 100 µg/mL BSA) and then resuspended in proximity ligation mix (1x T4 DNA ligase buffer, 100 µg/mL BSA, 20,000 U/mL T4 DNA ligase (New England Biolabs, #M0202S)). Proximity ligation was carried out by incubation overnight at RT with gentle mixing. Following proximity ligation, nuclei were pelleted and resuspended in a mix for removal of unligated DNA ends (1x NEBuffer 1 (New England Biolabs, #B7001S), 5 U/µL Exonuclease III (New England Biolabs, #M0206)) and then incubated with interval mixing for 15 minutes at 37 °C. Samples were reverse crosslinked by addition of 2 mg/mL Proteinase K, 1% SDS, 0.1 mg/mL RNase A, and 200 mM NaCl and overnight incubation at 65 °C with 1,000 rpm mixing. DNA was subsequently purified using the Zymo DCC-25 kit. Purified DNA was run on a 1% agarose gel to separate mono- and dinucleosomal bands. Dinucleosomal-sized fragments were excised from the gel and purified using the Zymoclean Gel DNA recovery kit (Zymo Research, #D4008). The purified biotinylated DNA was pulled down using MyOne Streptavidin T1 Dynabeads (Invitrogen, #65601) suspended in bead wash (BW) buffer (5 mM Tris-HCl pH 7.5, 1 M NaCl, 0.5 mM EDTA) at RT overnight. Beads were washed twice with Tween BW (TBW) buffer (0.1% Tween 20, 5 mM Tris-HCl pH 7.5, 1 M NaCl, 0.5 mM EDTA) and once with 10 mM Tris-HCl pH 7.5 before resuspending in 0.1x TE buffer. Micro-C libraries were prepared from the Dynabead-bound DNA using the NEBNext Ultra II DNA Library Prep kit (New England Biolabs, #E7645) with all incubations up to the end of adaptor ligation including interval mixing to maintain Dynabeads in suspension. After adaptor ligation and prior to PCR, beads were washed twice with TBW and once with 10 mM Tris-HCl pH 7.5. The number of cycles for PCR amplification were determined by running a test PCR on an aliquot of the main sample. Following PCR, Micro-C libraries were purified using 0.9x AmPure XP beads (Beckman Coulter, #A63880) according to the library prep kit protocol.

To capture reads originating from specific regions of interest, a custom probe panel (Twist Bioscience, #101001) was designed with 80-mer probes covering 14 genomic regions (see Table S1 for a list of genomic regions, and Table S2 for a list of probe locations (mm10 genome)). Regions were selected for the mm10 genome based on interactions between Polycomb loci visible in existing Micro-C datasets[25,26], and probes were filtered according to Twist's medium stringency filtering. This custom panel was used with the Twist Target Enrichment Standard Hybridization v2 Protocol to generate RCMC libraries. The number of PCR cycles to use for final RCMC library amplification was determined by running a test PCR on an aliquot of the main sample. RCMC library concentrations were determined by qPCR and sequenced (100 cycles, paired-end) on the NovaSeq X sequencing platform at the Broad Institute of MIT and Harvard.

#### ChIP-seq analysis

Native and crosslinked ChIP-seq data were processed the same way. fastq format reads were aligned to a concatenated version of the mm39 build of the *Mus musculus* genome and either the hg38 build of the *Homo sapiens* genome (crosslinked ChIP) or the dm6 build of the *Drosophila melanogaster* genome (native ChIP). Reads were aligned using Bowtie2[27] with the --no-mixed --no-discordant options. Multiply mapped reads were discarded by removing any reads containing the "XS" flag, and unmapped reads with the SAM flag 4 were also discarded. Duplicate reads were removed using sambamba markdup[28], and then split between reads aligning to mm39 or the spike-in genome. mm39-aligned reads in pulldown samples were downsampled according to the ratios of mm39 to spike-in reads in the pulldown and in the corresponding input for that sample, and normalized between all replicates and conditions for the same cell line.

bigWigs were generated from downsampled bam files using a custom Python script which uses MACS3 callpeak[29] with the -f BAMPE option, followed by wigToBigWig[30] from the UCSC suite to convert to bigWig format.

Peaks for RING1B and SUZ12 were taken from a previously published set (GSE119618)[21], and RING1B/SUZ12 shared peaks identified using bedtools[31] intersect with default options. The resulting overlapped peaks (Table S6) were converted to mm39 using UCSC LiftOver[32], giving a total of 5,627 peaks. RING1B/SUZ12 overlap peaks were used for all ChIP-seq metaplots and heatmaps with the exception of CBX2 ChIP in the $CBX2^{deg}$ cell line, where poor signal-to-noise ratio led to significant background. For metaplots for CBX2 ChIP in $CBX2^{deg}$, CBX2 peaks were called with a custom script using MACS3 with the -f BAMPE, --broad, -q 0.01 options and using the corresponding input from the higher quality CBX2 ChIP-seq for $PRC1/2^{deg}$ UNT and dTAG conditions. Peaks called in the dTAG condition were used to blacklist the peaks from the dTAG condition as non-specific peaks, and the peak set was manually curated to remove peaks from regions with poor mappability, leaving 875 peaks (Table S7).

Metaplots and heatmaps were generated using the above peaks from merged downsampled bigWigs with deepTools[33] computeMatrix in the reference-point mode with 10 kb flanks, using the --missingDataAsZero and --skipZeros options, followed by plotHeatmap.

#### $TT_{chem}$-seq analysis

$TT_{chem}$-seq data were aligned to a concatenated version of the mm39 build of the *Mus musculus* genome and the dm6 build of the *Drosophila melanogaster* genome. A custom Python script based on previous code[21] was used to first align reads to the rDNA sequences of both genomes using Bowtie2[27] with the --very-fast --no-mixed --no-discordant options. Unaligned (non-rRNA) reads were then aligned to the complete, concatenated genomes with STAR[34], and any reads with MAPQ values <4 were discarded. Any remaining unmapped reads were aligned again using Bowtie2 using the --sensitive-local --no-mixed --no-discordant --mm options. Multiply mapped reads were discarded by removing any reads containing the "XS" flag, and unmapped reads with the SAM flag 4 were also discarded. Aligned reads from STAR and Bowtie2 were merged and filtered for duplicates, and then split between mm39- and dm6-aligned reads. Aligned mm39 reads were downsampled between all replicates and conditions for a given cell line, normalizing to the number of reads arising from the mitochondrial genome to account for any nascent RNA labelling bias between samples.

bigWigs were generated from downsampled bam files using a custom Python script which splits reads according to their mapped strand using samtools and generates bigWigs for each using genomeCoverageBed from the bedtools[31] package. Merged bigWigs were generated for each condition using sambamba merge[28] to combine downsampled bam files.

Differentially expressed genes were determined using the DESeq2[35] R[36] package. First, counts were determined per gene using a previously described Perl script[21] (https://github.com/nFursova/Calibrated_ChIPseq_RNAseq/blob/main/RNAseq/RNAseq_GFF_PairedEnd_RNA_Read_Count.pl). The DESeq2 package was then used to process the read counts (DESeq2 outputs are available at GSE341537). Mitochondrial RNA normalization was applied by filtering for reads mapping to the mitochondrial genome and using these reads to calculate size factors for normalization. Differentially expressed genes were determined to be significant if the fold change was >1.5 and the p-adj was <0.05.

#### RCMC analysis

RCMC data were aligned and contact maps generated using a previously published Snakemake pipeline (https://github.com/ahansenlab/MicroC_RCMC_analysis)[24]. mcool files were balanced in the pipeline using a bed file of the genomic regions included in the custom probe panel to generate a blacklist for the rest of the genome to ignore reads in which at least one end did not map to one of the captured loci.

Polycomb-mediated interactions for aggregate pileup analysis were identified using the RING1B/SUZ12 overlap peaks as described in the crosslinked ChIP-seq section. Peaks within each of the captured regions were combined in a pairwise manner (n = 1,047). For each of the resulting pairs, observed over expected interaction strength was calculated in each replicate of each condition using a modification of a previously described analysis[23]. Any interactions where the interaction strength calculation failed due to a lack of data in any condition were discarded. Geometric means were then calculated for each interaction in each condition. The resulting interactions were then filtered according to the following metrics. First, interactions for which the standard deviation of the geometric mean of the untreated replicates was ≥1.3 were categorized as "noisy", due to high variability (n = 335). Next, interactions in which the shortest distance between the two anchors was less than the length of either anchor were removed ("short", n = 38), to remove very short loops between large Polycomb-occupied sites, such as *Hox* loci. Last, interactions for which the mean untreated interaction strength across all cell lines was less than 1.1 were removed, as "weak" interactions (n = 157). The remaining 517 interactions ("good") were used for downstream analysis. All pairwise combinations of sites with their categorizations can be found in Table S8.

Interactions between Polycomb-occupied sites were annotated as having CTCF at both anchors or not using a previously developed script (https://github.com/ahansenlab/RCMC_analysis_code/blob/main/loopFeatureOverlap.R)[23] using all interactions classified above as "good" and combined CTCF sites with SMC1 ChIP overlap generated previously[23].

Aggregate pileups were generated using the coolpup.py tool[37] with contact map binning at 5 kb. Expected values were computed for each chromosome with data, and pileups were generated with 50 kb flank sizes, rescaling interaction anchors to a third of the flank size and the pileup to 99 total pixels. Normalized 1D cross-sections of pileups were generated by averaging the 21 central diagonals from top right to bottom left of each pileup and normalizing all values to the mean of the 10 left-most and right-most (20 in total) values in the window.

RCMC data and all other genomics tracks were visualized for figure generation using CoolBox[38] within a custom Jupyter notebook.

#### Preparation of bead samples

To obtain the averaged point-spread function used in deconvolving Polycomb body images, a beads sample was prepared on 35 mm plates containing a 14 mm No. 1.5 coverglass (MatTek, #P35G-1.5-14-C). Wildtype mESCs were plated on a gelatinized MatTek dish and allowed to adhere overnight. The next day, cells were rinsed once with PBS, fixed for 15 minutes with 4% paraformaldehyde (ThermoFisher, #28908), then washed 4 times with PBS. Following the final wash, PBS was aspirated and 1 mL PBS containing 20 µL of 0.2 µm TetraSpeck microspheres (ThermoFisher, #T7280) sonicated for 5 minutes at room temperature with a VEVOR Ultrasonic Cleaner was added. Beads were allowed to adhere to the coverglass and fixed cell surfaces overnight at 4 °C, and stored in 2 mL PBS. For chromatic aberration correction and deconvolution in locus distance analysis, beads were prepared on an 8-well LabTek II chamber slide (ThermoFisher, #155409) using 10 µL TetraSpeck microspheres.

#### Polycomb body imaging

For imaging Polycomb bodies, cells containing RING1B-HaloTag were plated on gelatinized 35 mm plates containing a 14 mm No. 1.5 coverglass (MatTek, #P35G-1.5-14-C) the day before imaging, in medium containing 50 nM $JFX_{650}$[39]. 4 h prior to imaging, medium was replaced with fresh medium containing 50 nM $JFX_{650}$, with or without 100 nM dTAG-13. 1 h prior to imaging, media were again replaced with imaging medium (formulated the same as regular mESC medium with KnockOut DMEM replaced with high glucose DMEM without phenol red (ThermoFisher, #31053036) and $JFX_{650}$ and dTAG-13 as above.

Cells were imaged on an LSM 900 microscope with an Airyscan 2 detector (Zeiss) using Zen 3.0 (blue edition) software (Zeiss, v3.0.79.0006). Imaging was performed with cells in a humidified incubation chamber at 37 °C and 5% $CO_2$. For Polycomb body z-stacks, a Zeiss Plan-Apochromat 63x/NA1.40 objective was used with Immersol 518F immersion oil for 37 °C (Zeiss, #6S20-C0W9-F00N-01PF). For all cell lines, images were acquired from three separate treatments on different days. Six z-stacks were acquired on each day of imaging using the following acquisition parameters: 640 nm laser: 5% power, 780 V gain, 0 offset, 1.0 digital gain, 0.76 µs pixel time, with the emission filter set to permit wavelengths between 628-700 nm. Images were acquired using the SR-2Y acquisition mode with 200 nm z-spacing and 1.3x scan zoom, resulting in pixels measuring 55.9 nm in x and y, and a total frame size of 77.34 µm by 77.34 µm. For downstream processing, five z-stacks of TetraSpeck beads were taken using the same imaging settings with the exception of the 640 nm laser being set at 4% power.

#### Locus distance measurement imaging

For imaging of individual genomic loci, between 200,000 and 300,000 cells for each cell line were plated on two wells of an 8-well LabTek II chamber slide (ThermoFisher, #155409) the day before imaging. Cells were imaged in mESC growth media. For each cell line, one well was treated with 100 nM dTAG-13 for 4 hours and imaged within 4-6 hours of starting dTAG-13 treatment. Live-cell imaging of individual genomic loci was performed on a Zeiss Lattice Lightsheet 7 microscope equipped with 3 lasers (488 nm, 561 nm, 640 nm), two objectives (3.3x/0.4NA excitation objective, 44.83x/1.0 NA detection objective), and two Hamamatsu ORCA-Fusion sCMOS cameras using Zen 3.12/3.13. Sequential 561 nm and 488 nm illumination using two tracks on a single camera was used to collect dual-color images without camera offset. Acquisition was conducted using 20 ms exposure on the 15x650 lightsheet setting at 200 nm step size in the x dimension, with pixels within each sheet measuring 145 nm x 145 nm. The field of view was set to a height of 228 pixels, and the width and the number of slices for each stack were determined for each image based on the size of the mESC colonies being imaged. Between 3% and 10% power for 561 nm and between 20% and 40% power for 488 nm were used depending on the brightness of the colonies. Between 6 and 18 stacks were taken each day for each treatment replicate per cell line. At least 5 images of TetraSpeck beads were taken on each imaging day using 100% 561 nm and 100% 488 nm illumination.

#### Polycomb body image analysis

Acquired z-stacks were first Airyscan processed using Zen 3.0 software, with the 3D processing and Auto Filter settings, and at standard strength. Airyscan processed images were further processed using an adaptation of a previously published image analysis pipeline, Fyrtarn (https://github.com/ahansenlab/Fyrtarn)[12]. Images were first resampled to create isotropic pixel volumes of 55.9 nm in all x, y, and z dimensions. Resampled z-stacks were deconvolved using an implementation of Richardson-Lucy with Total-Variation regularization, using an averaged point spread function obtained from experimentally acquired bead images, and nuclei were segmented in 3D. Any nuclei touching the edges of the field of view were filtered out such that only nuclei entirely contained within the z-stacks were analysed. Subsequently, a subset of each nucleus was created from the resampled z-stack and the deconvolved z-stack, and Polycomb bodies were segmented using these subsets. Briefly, background subtraction was performed by applying 3D Gaussian blur to the resampled nucleus subset and subtracting the result from the deconvolved subset. An intensity threshold for Polycomb bodies was determined for each nuclear subset by calculating the mean of the 99$^{th}$ percentile of all nuclear pixel intensities and the pixel intensity value 3 standard deviations above the median. 3D particle segmentation was performed on deconvolved nuclear subsets after thresholding to identify individual Polycomb bodies. The coordinates, volumes, and mean pixel intensities of each Polycomb body, as well as the volume and mean intensity of the nucleus, were measured. All intensity measurements were obtained from the resampled z-stack. Manual quality check was performed by visually inspecting maximum intensity projections of each nucleus subset, and nuclei that were erroneously segmented (e.g. had extraneous background segmented with a nucleus, or two nuclei merged to one segmentation) were excluded from the final dataset to ensure accuracy of the nuclear volume. Outputs from nuclei and Polycomb body segmentation were analyzed using a custom R[36] script based on previous analysis[7]. For Polycomb bodies, segmented objects 27 voxels or smaller after isotropic resampling were discarded.

#### Locus distance analysis

3D distances between genomic loci were determined using a modification of the previously published Fyrtarn (https://github.com/ahansenlab/Fyrtarn)[12] pipeline. To run Fyrtarn on single snapshots rather than time-lapse data, start and end parameters were set to 0 and 1 respectively. Briefly, raw x-stacks from the Lattice Lightsheet were deskewed, coverglass transformed to yield isotropic 145 nm volumes in x, y, and z, then deconvolved using an implementation of Richardson-Lucy with Total-Variation regularization in Fyrtarn. Nuclei were segmented in 3D, and OR3 and synBsr1 foci were detected and their subpixel localization was determined using FracShift[40]. Foci on the same allele were paired based on a search distance of 5 pixels. Chromatic aberration correction was performed using TetraSpeck bead images, and 3D distances between all paired OR3 and synBsr1 foci were measured. Manual quality check was then performed to exclude all replicated sister chromatid foci and mis-identified foci, and only 3D distances between the focus pairs that passed the manual quality check were kept. Distributions of 3D distances were generated and plotted from exported data using a custom R script.

#### SUPPLEMENTARY TABLES

**Tables S1-S8.**

- **Table S1.** Chromosome positions of 14 genomic regions captured in RCMC provided with both mm10 and mm39 coordinates.
- **Table S2.** Chromosome locations of all regions covered by probes used in RCMC, given in mm10 coordinates.
- **Table S3.** Primer names and sequences for all cloning and screening primers used in this study.
- **Table S4.** sgRNA targets and sequences used for genome editing in this study.
- **Table S5.** Antibodies used for western blotting and ChIP-seq in this study.
- **Table S6.** Genomic loci with RING1B and SUZ12 ChIP-seq peak overlap based on peak calls from Fursova et al. 2019 (GSE119618).
- **Table S7.** CBX2 ChIP-seq peaks filtered for non-specific and low-mappability sites.
- **Table S8.** Genomic coordinates of Polycomb-Polycomb interactions categorized according to filtering for interaction strength, length, and noise.
